# Group composition and dynamics in American Crows: insights into an unusual cooperative breeder

**DOI:** 10.1101/2024.02.01.578454

**Authors:** Carolee Caffrey, Charles C. Peterson

**Affiliations:** Zoology Department, Oklahoma State University, Stillwater OK 74075, USA; 5235 Kester Ave #318, Sherman Oaks CA, USA; Copper Mountain College, Joshua Tree CA, 92252, USA

## Abstract

Breeding pairs of American Crows (*Corvus brachyrhynchos*) in Stillwater, OK, lived with 0-10 auxiliaries in territories distributed throughout public, campus, commercial, and residential areas. Unpaired crows moved easily among groups throughout the year, but commonly did so during the two months or so preceding the onset of nesting across the population, and the week or so preceding hatching within groups. In 2001 and 2002, pre-hatch group size ranged from 2-10 (mean = 4.5 in both years), and auxiliaries included a male sibling, social and genetic offspring, step-offspring, half-siblings, and unrelated immigrants of both sexes, ranging in age from 1 to at least five years old. Twenty nine percent of pre-hatch auxiliaries dispersed out of groups at hatching (for half, only temporarily), including all females unrelated to female breeders. Post-hatch group size ranged from 2-6, with means of 3.7 in both years, and the post-hatch auxiliary population differed in composition from the pre-hatch population: whereas post-hatch male auxiliaries included a sibling, half siblings, and unrelated immigrants in addition to social and genetic sons, all post-hatch female auxiliaries were the social and genetic daughters of female breeders, and all but one (the same individual in both years) were also the social and genetic daughters of male breeders.

Crows in Stillwater delayed breeding for one or more years beyond sexual maturity, despite the availability of space and members of the opposite sex. Individual dispersal decisions by unpaired crows, and the behavior of paired territory owners, did not follow patterns described for other cooperative breeders. We found little support for extant theories regarding the formation, composition, and maintenance of avian groups, and discuss aspects of the lives of crows that may have contributed to the complex and benign nature of this population’s society.

How to Cite: Caffrey, C. and C. C. Peterson. 2015. Group composition and dynamics in American

Crows: insights into an unusual cooperative breeder. Friesen Press.

## INTRODUCTION

Published ideas on possible reasons for the formation of groups in cooperatively breeding birds date back to at least the 1960s: Selander (1964, in Brown 1978 and Koening et al. 1992) and Brown (1969) both hypothesized that crowded breeding conditions might underlie the delay of independent reproductive attempts by mature individuals, who possibly stood to gain higher benefits over the long run by currently remaining in natal territories. For many years thereafter, explanations for group formation tended to develop in association with what came to be the “ecological constraints hypothesis” (formalized by Emlen 1982, 1984): that prohibitive environmental factors rendered natal dispersal and independent breeding disadvantageous, and so newly mature individuals were selectively forced to forego breeding and remain in natal territories (often then, and sometimes still, discussed in the literature as if the two were a single decision). As ideas continued to expand, potential direct fitness advantages associated with delaying dispersal came to be recognized and formalized as the “benefits-of-philopatry hypothesis” (Stacey and Ligon 1987, 1991, Zack 1990). Although a merger of the two sides of the “semantic coin” (Emlen 1995) was under-way years earlier (e.g., Brown 1974), it wasn’t until the early 1990s that it came to be widely understood and acknowledged that unpaired individuals will assess the consequences of different behavioral options, in contexts of current situations, and subsequently behave in ways that maximize their lifetime fitness (Walters 1991, Emlen 1991, 1994, Koening et al. 1992, Komdeur 1992); individuals should disperse and attempt to breed when opportunities of worth are available, and if no such opportunities exist, individuals should pursue current next-best alternatives.

Given decisions to delay independent breeding, subsequent decisions to delay dispersal (as opposed to other options; below) – and so to reside in natal territories – have been hypothesized to be tipped by the many possible benefits to doing so, including increased survivorship (Ekman et al. 2004, Pruett-Jones 2004, and references therein) and the exploitation of opportunities to enhance future reproductive success (e.g., through territorial inheritance or budding, and using natal territories as safe “havens” [Ekman et al. 1999, Kokko and Ekman 2002] from which to monitor and successfully compete for available within-population vacancies [Koening et al. 1992, Emlen 1994, Cockburn 1998, and Ekman et al. 2004, Dickinson and Hatchwell 2004, Pruett-Jones 2004 and references therein]), as well as benefits derived through nepotism (access to parental resources and possible assistance in securing breeding status by genetic relatives [Ekman et al. 2001, 2004]) and group augmentation (Woolfenden 1975, Ligon and Ligon 1978, Wiley and Rabenold 1984, Clutton-Brock 2002, Dickinson and Hatchwell 2004, Bergmüller et al. 2007, and Kokko et al. 2001 and Stevens and Gilby 2004 and references therein).

Cost/benefit treatments in the cooperative-breeding literature generally revolve around whether or not individuals should disperse and attempt to breed, generally involve weighing the possible direct benefits of independent breeding against benefits postulated for delaying dispersal and “helping” (the latter two also too often discussed as if they were a single decision), and have largely been applied only to species in which sexual maturation occurs in an individual’s second year (e.g., Dickinson and Hatchwell 2004, Ekman et al. 2004). Individuals that delay dispersal tend to be thought of as “sedentary” (e.g., Ekman et al. 2004). Heuristic comparisons are often among groups of different sizes and composition, without much consideration of the day-to-day interactions among individuals that likely influence the fitness outcomes of different dispersal choices (Wilson 1998, Reale et al. 2007, Taborsky and Oliveira 2012). Rarely discussed in the cooperative-breeding literature are alternatives to delaying dispersal for individuals as independence is achieved, or for unpaired group members as nesting seasons approach, although “floating” has received some attention, particularly recently (e.g., Brown 1969, Verbeek 1973, Smith 1978, Koening et al. 1992, Zach and Stutchbury 1992, Bruinzeel and van de Pol 2004, Penteriani et al. 2011, Lenda et al. 2012, Penteriani and Delgado 2012, Moulton et al. 2013).

Diversity in breeding-population social organization is characteristic of the Clade Corvida (Cockburn 1996), the Family Corvidae (Ekman et al. 2004), and the genus *Corvus*. Pairs of Common Ravens (*C. corax*; Boarman and Heinrich 1999, Webb et al. 2012), Chihuahuan Ravens (*C. crytoleucus*; Bednarz and Raitt 2002, D’Auria and Caccamise 2007), Fish Crows (*C. ossifragus*; McGowan 2001a), and Mariana Crows (*C. kubaryi;* S Faegre, pers. comm.) almost always breed socially monogamously, as do pairs of Carrion Crows (*C. corone)* in most populations (references in Baglione et al. 2002a and 2005), yet many pairs in at least one population of Carrion Crows breed cooperatively in groups that include up to seven auxiliaries (Baglione et al. 2002a). Pairs of Northwestern Crows (*C. caurinus*; Verbeek and Butler 1999) and New Caledonian Crows (*C. moneduloides*; Holzhaider et al. 2011) breed socially monogamously or unassisted in small groups, pairs of American Crows (*C. brachyrhynchos*; Chamberlain-Auger et al. 1990, McGowan 2001b, Verbeek and Caffrey 2002) breed socially monogamously or cooperatively in groups including up to 10 auxiliaries, and pairs of Rooks (*C. frugilegus*; Coombs 1978), Jackdaws (*C.monedula*; Henderson et al. 2000), and Western American Crows (C*. b. hesperis*; Caffrey 1992, 2000) nest colonially (Rooks and Jackdaws always without assistance, Western American Crows sometimes with assistance). Nonbreeding individuals are known to group into aggregations during breeding seasons in three populations with different social structure: loose groups of nonbreeders spread across large areas in Common Ravens (Braun and Bugnyar 2012), and resident flocks amid populations of Carrion Crows (Baglione et al. 2005) and Western American Crows (Caffrey 1992). Information is available in the literature on group size and composition for populations of Western American Crows, Carrion Crows, American Crows, New Caledonian Crows, and Mariana Crows (see Discussion, Intra- and inter-specific comparisons, below), but nowhere exist details of the relationships among individuals that manifest as the population and group characteristics researchers have measured. To wit, in the words of Clayton and Emery (2007), we are “completely ignorant of the social lives of most of the extant corvids.”

Here, we document the unusual nesting-season social organization of a population of American Crows in Stillwater, OK. Members of this population lived as pairs or in groups of up to 12 individuals (Verbeek and Caffrey 2002) that bred cooperatively. Crows are unlike many other cooperative breeders in that individuals do not mature sexually until their third year (Black 1941, Townsend et al. 2009). Also unlike (in) many other cooperative breeders, the social organization in our study population was dynamic, in that unpaired individuals of both sexes and diverse ages moved among groups throughout the year (CC, unpubl. data). We examined the genetic and social relationships among population members in search of support for classic and current theories on the formation of groups in cooper-atively-breeding birds (above), and address the need for continued expansion of such ideas, at least as they apply to crows.

## METHODS

This study was carried out in strict accordance with the recommendations in the Guidelines to the use of Wild Birds in Research of The Ornithological Council, and was approved by the Oklahoma State University IACUC (ACUP No: AS50713). Capture and marking of crows occurred under U.S. Department of Interior, U.S. Geological Society, Federal Bird Banding Permit #22165 (Caffrey).

### Background

-Our study population of marked crows in Stillwater, Oklahoma, had been under observation since August 1997, yet it took several years to get a majority of population members marked. For the present study, crows were observed during the nesting seasons (including nest building, incubation, and nestling care stages) of 2001 and 2002, late February through mid- and early May, respectively, in the two years. During those years, Stillwater was a university town of approximately 40,000 people within a 72.51 km^2^ area (U.S. Census Bureau). Crows in Stillwater based their activities in all-purpose territories throughout the year; territories ranged in size from approximately 50 to 150 ha and were distributed throughout public, campus, commercial, and residential areas (Fig. 1). Observations were made with binoculars and spotting scopes, primarily from vehicles, except for one group in 2001 (#29: Tables 1 and 3, and Appendix A 1a), whose placement of their nest next to a window in an empty building allowed occasional continuous videotaping of nest activities for up to 10 hours at a time.

**FIG. 1.**
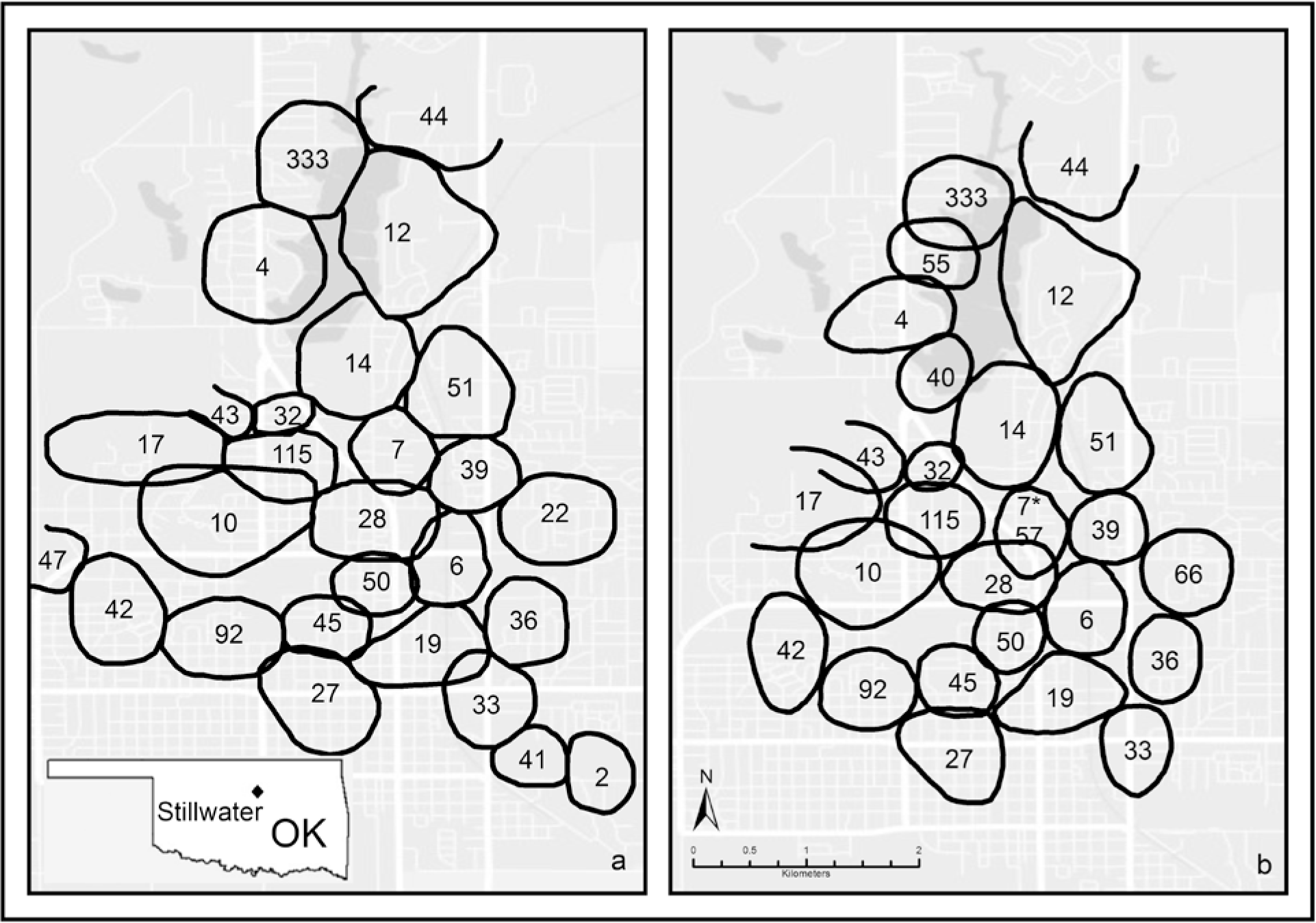
Approximation of the territories of study groups of crows in Stillwater OK in a) 2001, and b) 2002. In 2002, Group 7 changed to Group 57 upon the replacement of the female breeder (Appendix A 2c).

**Table 1.**
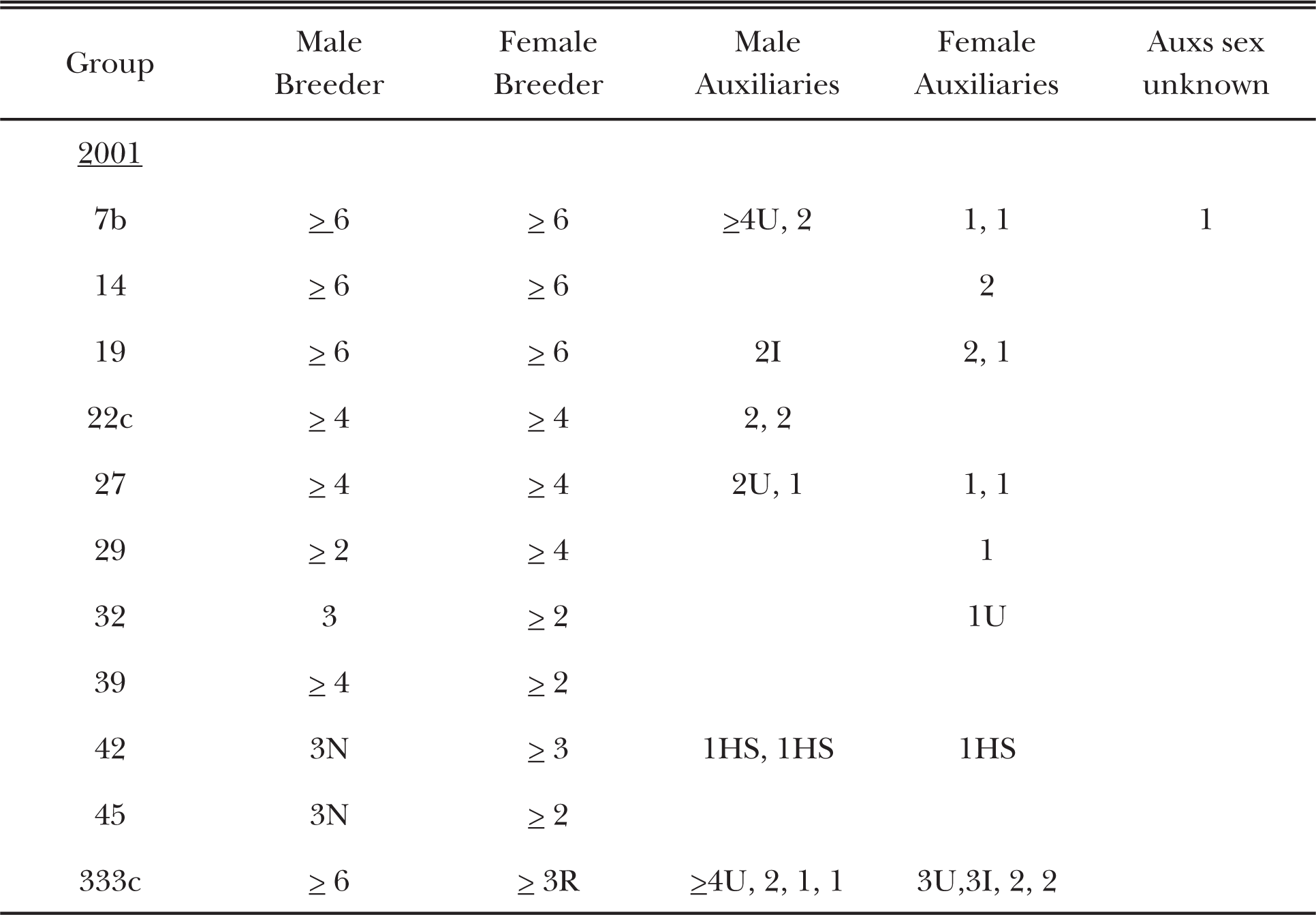
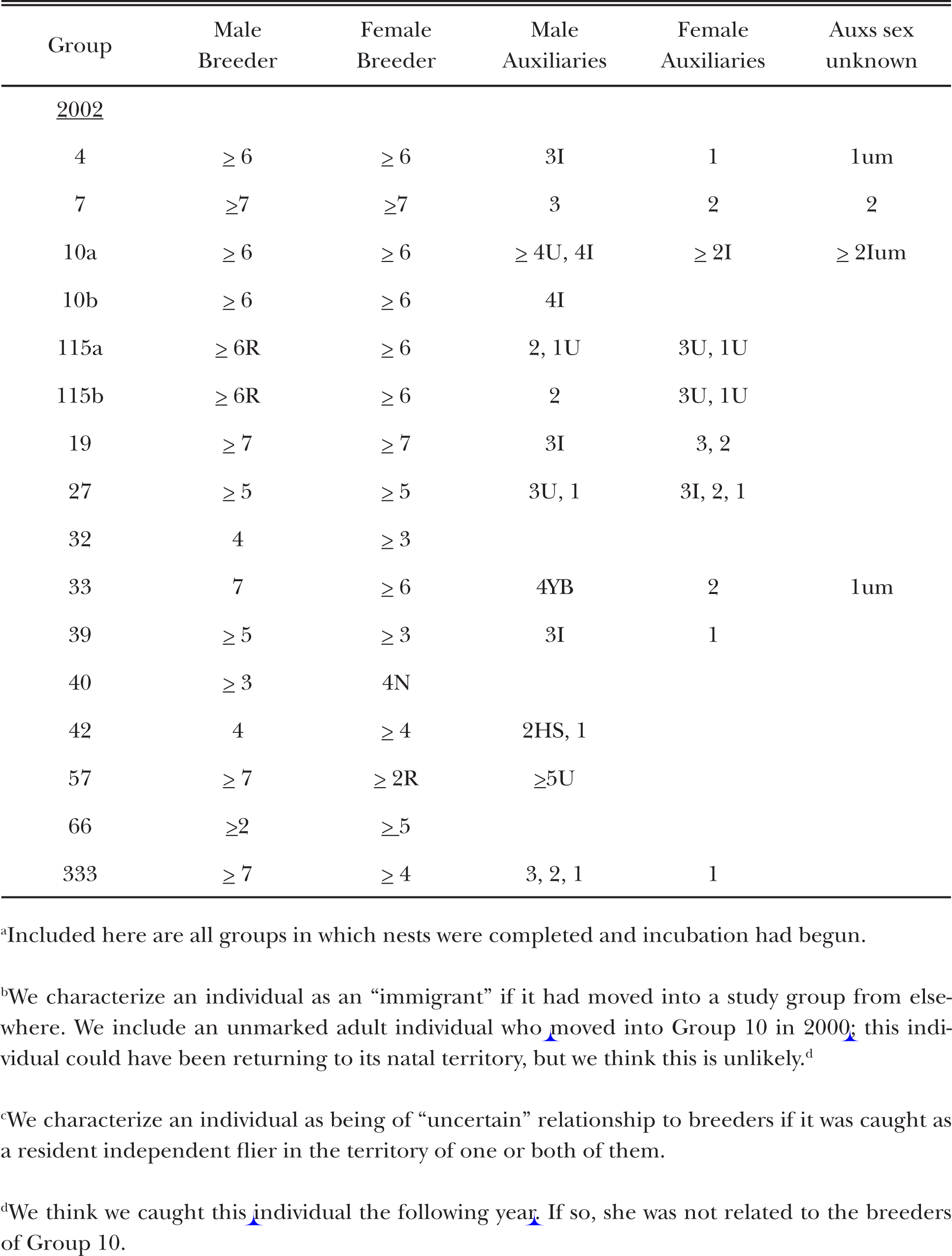
Composition of pre-hatch crow groups^a^. Each numeral represents an individual by its age in years. Most auxiliaries had hatched in nests of one or both breeders. N (novice): known to be breeding for first time. R: breeder was replacement of social parent of at least one of auxiliaries in group. I: immigrant^b^. HS: half-sib of male breeder. YB: younger brother of male breeder. U: dispersal history (relative to group) uncertain^c^. um: unmarked.

| Group | Male Breeder | Female Breeder | Male Auxiliaries | Female Auxiliaries | Auxs sex unknown |
| --- | --- | --- | --- | --- | --- |
| <u>2001</u> |  |  |  |  |  |
| 7b | ≥ 6 | ≥ 6 | ≥4U, 2 | 1, 1 | 1 |
| 14 | ≥ 6 | ≥ 6 |  | 2 |  |
| 19 | ≥ 6 | ≥ 6 | 2I | 2, 1 |  |
| 22c | ≥ 4 | ≥ 4 | 2, 2 |  |  |
| 27 | ≥ 4 | ≥ 4 | 2U, 1 | 1, 1 |  |
| 29 | ≥ 2 | ≥ 4 |  | 1 |  |
| 32 | 3 | ≥ 2 |  | 1U |  |
| 39 | ≥ 4 | ≥ 2 |  |  |  |
| 42 | 3N | ≥ 3 | 1HS, 1HS | 1HS |  |
| 45 | 3N | ≥ 2 |  |  |  |
| 333c | ≥ 6 | ≥ 3R | ≥4U, 2, 1, 1 | 3U,3I, 2, 2 |  |
| <u>2002</u> |  |  |  |  |  |
| 4 | ≥ 6 | ≥ 6 | 3I | 1 | 1um |
| 7 | ≥ 7 | ≥ 7 | 3 | 2 | 2 |
| 10a | ≥ 6 | ≥ 6 | ≥ 4U, 4I | ≥ 2I | ≥ 2Ium |
| 10b | ≥ 6 | ≥ 6 | 4I |  |  |
| 115a | ≥ 6R | ≥ 6 | 2, 1U | 3U, 1U |  |
| 115b | ≥ 6R | ≥ 6 | 2 | 3U, 1U |  |
| 19 | ≥ 7 | ≥ 7 | 3I | 3, 2 |  |
| 27 | ≥ 5 | ≥ 5 | 3U, 1 | 3I, 2, 1 |  |
| 32 | 4 | ≥ 3 |  |  |  |
| 33 | 7 | ≥ 6 | 4YB | 2 | 1um |
| 39 | ≥ 5 | ≥ 3 | 3I | 1 |  |
| 40 | ≥ 3 | 4N |  |  |  |
| 42 | 4 | ≥ 4 | 2HS, 1 |  |  |
| 57 | ≥ 7 | ≥ 2R | ≥ 5U |  |  |
| 66 | ≥ 2 | ≥ 5 |  |  |  |
| 333 | ≥ 7 | ≥ 4 | 3, 2, 1 | 1 |  |

<sup>a</sup>Included here are all groups in which nests were completed and incubation had begun.
<sup>b</sup>We characterize an individual as an “immigrant” if it had moved into a study group from elsewhere. We include an unmarked adult individual who moved into Group 10 in 2000; this individual could have been returning to its natal territory, but we think this is unlikely.<sup>d</sup>
<sup>c</sup>We characterize an individual as being of “uncertain” relationship to breeders if it was caught as a resident independent flier in the territory of one or both of them.
<sup>d</sup>We think we caught this individual the following year. If so, she was not related to the breeders of Group 10.

We captured first-calendar-year, second-year, and adult crows with a rocket net and a Netlauncher (Caffrey 2002a). Sexually immature, second-year crows (“yearlings” or “1-year olds” hereafter) were distinguishable from older, adult individuals via use of plumage and mouth-color characteristics (Emlen 1936). Nestlings were temporarily removed from nests prior to fledging for processing (Caffrey 2002b and c). We marked all crows with patagial tags bearing two-letter names (Appendix D) and unique combinations of colored plastic and U.S. Fish and Wildlife Service metal leg bands (Caffrey 2002b). No group contained more than one unmarked individual. Because the actual ages of crows caught as adults could not be determined, we assigned them the age “≥2 y” at marking. We caught many breeders as members of already-established pairs, and because all but one crow (of known age) in our study population did not breed until at least three years old (CC, unpubl. data), many were likely older than the minimum ages reported in Table 1. Pairs of crows sometimes renested after failed attempts; we identify first, second, and third attempts (ones wherein incubation of eggs had begun) per season as “a,” “b,” and “c.”

We determined the timing of the onset of incubation mostly through observation of female breeder behavior (Caffrey et al. 2015b). Hatch dates were determined primarily via nest watches and were identified as the days feeding trips were first observed (Caffrey et al. 2015a).

### Group Composition

-During this study, groups included from 0-8 “auxiliaries” (defined below). We identified as breeders pairs of adult females and males most active in territory defense, courtship, and nest building.

Because a common time for group members to disperse out of groups was during the week or so preceding hatching, we report separate data for pre-hatch and post-hatch crow groups. We used the behavior of the crows themselves with and toward each other, and the locations of their interactions, to identify groups and group members. As such, we defined as pre-hatch auxiliaries individuals seen with one or both breeders, during March and through incubation, on at least 20% of the occasions that at least one breeder was observed with non-pair group members. We defined as post-hatch auxiliaries individuals observed with at least one breeder in at least 35% of all observations of breeders away from nest areas. In Tables 1 and 3, we distinguish between auxiliaries hatched in nests of one or both breeders, auxiliaries that immigrated into study groups (did not reside with them previously during our study), and auxiliaries originally caught (and marked) as independent fliers in territories of study groups (and so whose dispersal histories, relative to breeders, were uncertain). We later determined parentage for eight of the nine individuals of uncertain relationship to breeders in Table 1 (= 13 auxiliaries; we lacked a DNA sample for one individual) and six of the seven in Table 3 (= eight auxiliaries); see below. Hereafter we use such terms as “social mother” and “social son” for relationships of nestlings and their attending breeders, and “daughter,” “brother,” “mother,” and “offspring” when the genetic parentage of particular individuals is known.

### Genetic Sampling and Analyses

-We collected approximately 1cc of blood from the brachial vein of each crow for molecular determination of sex and for estimation of microsatellite-based relatedness values. DNAs were extracted from blood samples following the method of Longmire et al. (1997). We sexed all individuals at diagnostic sex-linked alleles, using the 2550/2718 primer set (Fridolfsson and Ellegren 1999). For this study, we genotyped 151 individuals, including 63 nestlings (marked in 2001 and 2002) and 88 auxiliaries, social parents, and potential extrapair males, at 9-10 microsatellite loci, following Townsend et al. (2009). Genotyping was performed on a 3100 Genetic Analyzer (Applied Biosystems). All alleles were scored automatically and con-firmed visually using GENEMAPPER^TM^ version 3.7 software (Applied Biosystems).

We employed the maximum likelihood approach implemented in the program CERVUS 3.0 (Kalinowski et al. 2007). We specified a potential typing error of 1%, and specified the proportion of sampled candidate parents at 90% to account for unsampled adults in areas adjacent to our focal territories. We specified relatedness among 5% of candidate parents at 0.5 to account for kinship of potential breeders within family groups. These ten loci provided a powerful marker set for parentage discrimination, with a mean allele frequency of 13.1 alleles per locus and a combined non-exclusion probability of 0.00067000 for the first parent and 0.00001227 for the second parent. Allele frequencies at these ten loci did not deviate significantly from Hardy-Weinberg expectations.

We first tested the assumption that social mothers were the genetic mothers of all nestlings in their broods by specifying social mothers as the “known parent” in CERVUS 3.0 and examining their pairwise-match. Social mothers in this sample had zero (*n* = 50 pairs) to one (*n* = 20 pairs) mismatches with their putative offspring, the latter of which we attributed to genotyping error or mutation (Slate et al. 2000). We therefore accepted all social mothers as genetic mothers in this sample.

We then examined paternity, specifying female breeders as “known parents,” and including all sampled adult males present in a given year as potential fathers. Confidence levels of 80% or greater might be sufficient to identify true genetic parents when combined with behavioral data (Slate et al. 2000). We therefore accepted males suggested by CERVUS 3.0 as true sires when they were selected as the most likely candidate at or above the 80% trio confidence level in combination with the known mother. When the trio confidence level for the suggested candidate fell below 80%, we denoted those offspring as having sires of unknown identity (n = 2 nestlings).

We used our microsatellite genotypes to generate pairwise genetic relatedness coefficients (r) between all pairs of group members using the program RELATEDNESS v.5.0.8 (Queller and Goodnight 1989). Negative coefficients suggest that two individuals are less related than expected by chance if the two genotypes were randomly selected, whereas positive coefficients suggest that the individuals are related (e.g., mean coefficients of 0.5, 0.25, and 0.125 are expected between first-, second- and third-order kin, respectively). In accordance with these expectations, the mean coefficient of relatedness found in a preliminary analysis of 34 mother-offspring (first-order) kin pairs was 0.47 ± 0.02 SE (range = 0.11-0.72; 95% CI = 0.43-0.52). All pairs with relatedness coefficients at or below zero are likely to be non-relatives, and we therefore truncated the lower bound of the distribution at zero.

We did not have DNA samples for one adult female (pre- and post-hatch) auxiliary in 2001 and five auxiliaries (four individuals; one in both years) whose sex was unknown. Throughout this paper, where appropriate, we report r values as means ± 95% CIs.

## RESULTS

### Pre-hatch Group Composition

-We report the composition of pre-hatch groups during 11 nesting attempts in 2001 (including a second and two third attempts by three groups) and 16 in 2002 (including the first and second attempts of one group). Pre-hatch group composition was highly variable (Table 1), and intra- and inter-group social dynamics were complex (examples in Appendix A 1a and b).

In 2001 and 2002, 82 and 81% of pre-hatch groups, respectively, were pairs with at least one auxiliary. Group size ranged from 2-10 with means of 4.5 ± 2.4 SD and 4.4 ± 1.6 SD in the two years, and auxiliaries ranged in age from 1 to at least five years old (Table 1). The 66 total auxiliaries in pre-hatch groups in 2001 and 2002 included 48 individuals (22 males, 22 females, and four of unknown sex) of which 13 (7 males, 5 females, and one of unknown sex) were auxiliaries in both years (1 male and 1 female were each in different groups in the two years). There was no sex bias to the auxiliary population in either year (Wilcoxon Signed Ranks Test, p>0.35 in both years) and female and male auxiliaries differed in age only in 2002: in 2001, mean age of females = 1.6 ± 0.8 SD, of males = 1.9 ± 1.0 SD; in 2002, mean age of females = 1.9 ± 0.8 SD, of males = 2.7 ± 1.2 SD yrs (t33 = 2.29, p=0.029 [Table 1]).

Most auxiliaries in pre-hatch groups had delayed dispersal: of 51 of the 66 pre-hatch auxiliaries with known histories (Table 1), 36 (70%; 27 individuals) were previous nestlings of at least one breeder, and 30 (59%; 23 individuals) had been nestlings of both. Six auxiliaries (five individuals; one in attempts a and b in 2002) were in natal territories with a replacement for one of their parents, and four (three individuals; one in both years) were home with an older brother (NK, in Appendix A 1a) who had taken over ownership of his natal territory from his father. (It is also likely that the majority of the “Uncertains” [13 auxiliaries/9 individuals] in Table 1 had also delayed dispersal [below; parentage data].) Among auxiliaries with known histories for whom we had DNA, the mean related coefficient (r ± 95% CI) to social mothers in 2001 was 0.53±0.07 (n=14 auxiliaries), and 0.37±0.09 (n=17) to social fathers; in 2002, mean r to social mothers = 0.51±0.04 (n=17), to social fathers = 0.42±0.09 (n=16).

Of auxiliaries that had not delayed dispersal, one had moved with his older brother (RN) to a neighboring territory when RN paired with the (presumably widowed) female (Group 33; Appendix A 2a), and 10 were immigrants (nine individuals [one in both years]: mean r to female and male breeder in 2001 [n = 2 auxiliaries] = 0.09 and 0.0, respectively, and 0.01+0.02 and 0.04+0.06 in 2002 [n=7]). Thirteen pre-hatch auxiliaries (nine individuals, four in both years) had been caught as independent fliers with their groups and so their dispersal histories were uncertain. We later determined parentage for eight of these individuals: five were the offspring of both current breeders, one (FX; Appendix A 2b) was the daughter of the female breeder and unrelated to her mother’s current mate (a replacement for her father), one (PH; Appendix A 2c) was the son of the male breeder and unrelated to his father’s current mate (a replacement for his mother), and one (NX) was *not* the son of the breeders in the group in which he was caught (in 1999, but their microsatellite-based relatedness coefficient suggested he was not unrelated to the male [SW, in Appendix A 1b]). (In 2001 and early 2002, NX was in the same territory with SW but a different female breeder [DZ, to whom his microsatellite-based relatedness coefficient suggested he was also not unrelated; Appendix A 1b].) In addition to the above, in 2002, two groups each included an unmarked yearling, likely offspring from 2001 that had delayed dispersal. Overall, there were no differences in genetic relatedness of auxiliaries to the female and male breeders with which they resided through the building of nests and incubation of eggs (Fig. 2, left-most columns).

**FIG. 2.**
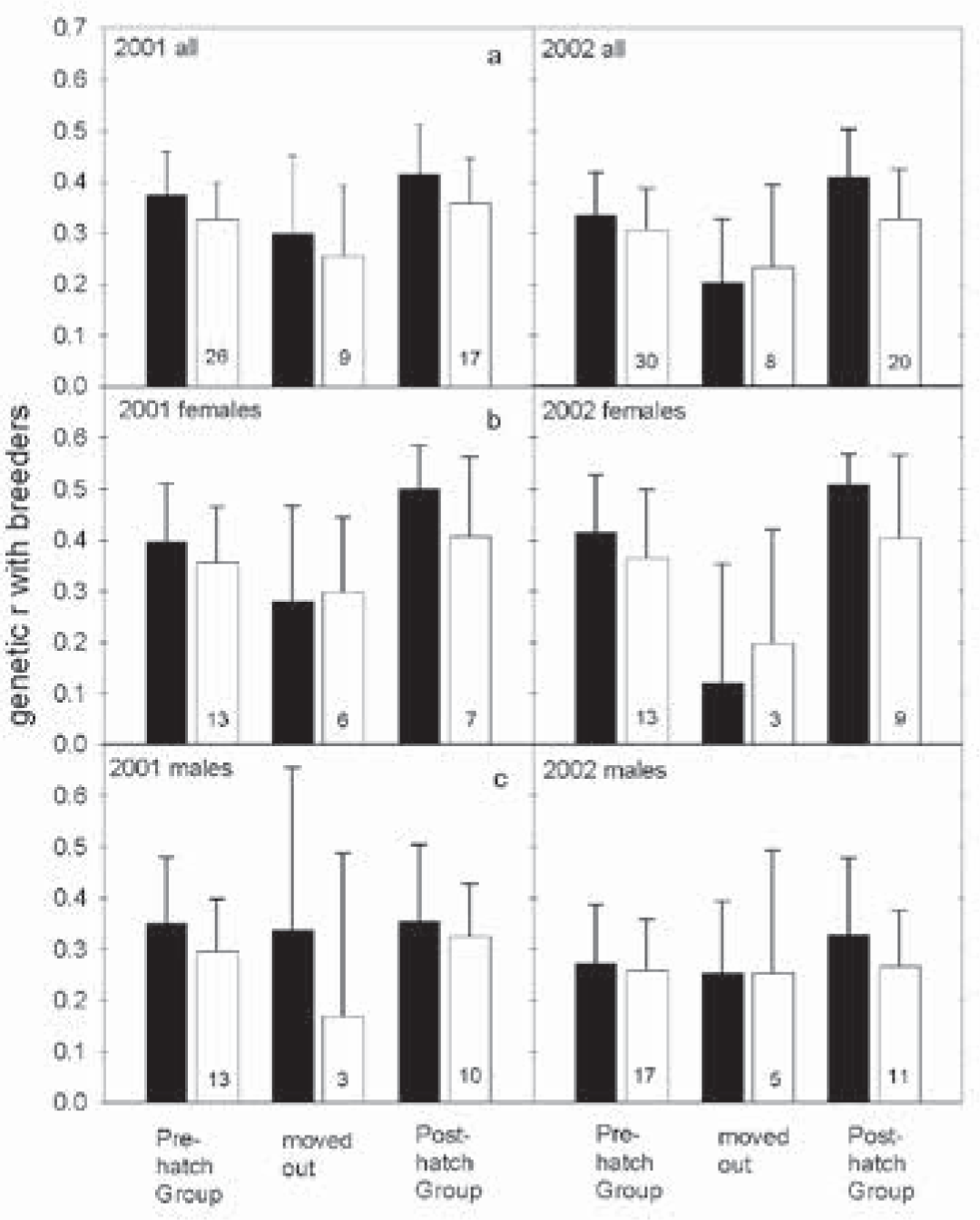
Mean (± 95%CI) genetic relatedness to breeders of auxiliary members of pre-hatch groups, pre-hatch auxiliaries that left groups, and post-hatch auxiliaries. Solid bars depict relatedness to female breeders; unfilled bars that to male breeders. Numerals indicate sample sizes.

Most immigrants had moved into study groups as adults: one (unmarked) individual of unknown sex and origin, three of three females (one from one territory away [PR; Appendix A 1b], one from 5-6 territories away [LH; Appendix A 1b and 3c], and one whose departure site was unknown), and two of four males. Two males moved into groups during their second year (one in April [TM; Appendix A 1a and 2d] and one in October [RG; Appendix A 1b]). All male immigrants in pre-hatch groups in 2001 and 2002 had moved from adjacent groups (not always the case in previous years [CC, unpubl. data]).

### Dispersal, or not, out of Pre-hatch Groups

-Table 2 tabulates the subsequent residency decisions of 62 pre-hatch auxiliaries (in groups where eggs hatched): most (44; 71%) chose to remain for nestling periods. Of the 18 (29% of pre-hatch auxiliaries) that dispersed out of groups during nesting seasons in the two years, 14 did so during the period from 10 days preceding to two days after hatching; the four exceptions included an adult immigrant female (LH) who left her group shortly after incubation began, possibly in response to harassment (Group 27 in 2002; Appendix A 1b), and three adult auxiliaries from the same group (#10, in 2002) that dispersed upon the breeding pair’s usurpation of the nest of one of the three dispersers (an unmarked individual) and a fourth auxiliary (TM; Appendix A 2d). The breeding pair and TM completed his (usurped) nest and raised three fledglings (Appendix A 2d). Although individual patterns of dispersal were variable, the majority of the 17 marked dispersers maintained contact with the groups from which they dispersed, and many moved back in with those groups at later dates (Appendix A 3).

**Table 2.** Within-nesting-season residency decisions of 62 pre-hatch auxiliary crows in 2001 and 2002^a^. Numerals indicate the number of individuals in each category.

| age | males |  | females |  | sex unknown |  |
| --- | --- | --- | --- | --- | --- | --- |
|  | Pre-Hatch | Post-hatch | Pre-hatch | Post-hatch | Pre-hatch | Post-hatch |
| <u>2001</u> |  |  |  |  |  |  |
| 1 | 5 | 4 | 8 | 5 | 1 | 1 |
| 2 | 6 | 4 | 4 | 2 |  |  |
| 3 |  |  | 2 | 1 |  |  |
| ≥4 | 2 | 2 |  |  |  |  |
| <u>2002</u> |  |  |  |  |  |  |
| 1 | 4 | 3 | 6 | 5 | 2 | 2 |
| 2 | 4 | 4 | 4 | 3 | 1 |  |
| 3 | 5 | 2 | 4 | 3 |  |  |
| 4 | 3 | 2 |  |  |  |  |
| ≥5 | 1 | 1 |  |  |  |  |

<sup>a</sup>Included are all individuals from all attempts of all groups with pre-hatch group data available, in which eggs hatched.

Although the relationships with breeders of auxiliaries that chose to leave and auxiliaries that chose to stay were variable, dispersers (including social and genetic daughters and sons of both breeders [n = 7], social daughters and sons of male breeders and their replacement mates [n = 4], a half-sib of the male breeder, and five immigrants) tended to be less closely related to breeders than were auxiliaries that remained for nestling periods, particularly in the case of female auxiliaries and female breeders (Fig. 2): all female immigrants, and female auxiliaries in groups with replacement female breeders, left, such that by two days after hatching had begun within groups, all female auxiliaries unrelated to female breeders were gone.

The three longest-distance dispersal events of which we are aware all involved females. One, a one-year old female (AI, the half-sister of the male breeder [Group 42, 2001; Table 1, and Appendix A 1a and 3f3]), present through nest building and most of incubation, made a brief foray for a few days to Pawnee, OK (44.9 km away, and the destination to which she would ultimately disperse), 10-14 days before hatching. She returned to her group temporarily and was observed to make one nestling feeding trip the day after hatching began before she moved out (in mid April). By late May (and thereafter) she was a permanent resident of Pawnee. Another, in her second year, was twice seen in Stroud, OK (a distance of 83 km), and the other, an adult, was twice seen in Norman, OK (136 km).

### Post-hatch Group Composition

-Post-hatch group size ranged from 2-6, with means of 3.7 ± 1.3 SD in 2001 (11 groups) and 3.7 ± 1.2 SD in 2002 (13 groups), auxiliaries ranged in age from one to at least five years old, and the composition of groups was variable (Table 3). In 2001, 77% of post-hatch groups were pairs with at least one auxiliary; in 2002, 82% were so. There was no sex bias to the post-hatch auxiliary population, or difference in the ages of female and male post-hatch auxiliaries, in either year.

**Table 3.** Composition of post-hatch crow groups. For breeders, experience: 1 = experienced/ nested with current mate in the previous year, 2 = experienced/did not nest with current mate in previous year (e.g., #66 in 2002) or attempt (#57 in 2002; Appendix A 2c), 3 = nesting for first time, 4 = likely nesting for first time, 5 = history unknown. For auxiliaries, each numeral represents an individual by its age in years. Most auxiliaries had hatched in nests of one or both breeders. I: immigrant^a^. HS: half-sib of male breeder. YB: younger brother of male breeder. U: dispersal history (relative to group) uncertain^b^. um: unmarked. x: disappeared from post-hatch group.

| Group | Male Experience | Female Experience | Male Auxiliaries | Female Auxiliaries | Sex unk Auxiliaries |
| --- | --- | --- | --- | --- | --- |
| <u>2001</u> |  |  |  |  |  |
| 7b | 1 | 1 | ≥4U, 2 |  | 1 |
| 14 | 1 | 1 |  | 2 |  |
| 19 | 1 | 1 | 2I | 2, 1 |  |
| 22c | 1x | 1 | 2 |  |  |
| 27 | 1 | 1 | 2U | 1, 1 |  |
| 29 | 1x | 1 |  | 1x |  |
| 32 | 1 | 1 |  | 1U |  |
| 39 | 4 | 5 |  |  |  |
| 42 | 3 | 5 | 1HS, 1HS |  |  |
| 45 | 3 | 5 |  |  |  |
| 333c | 1 | 1 | ≥4U, 1, 1 | 3Ux |  |

| Group | Male<br>Experience | Female<br>Experience | Male<br>Auxiliaries | Female<br>Auxiliaries | Sex unk<br>Auxiliaries |
| --- | --- | --- | --- | --- | --- |
| <u>2002</u> |  |  |  |  |  |
| 4 | 1 | 1 |  | 1 | 1um |
| 10b | 1 | 1 | 4I |  |  |
| 115b | 1 | 1 | 2 | 3U, 1 |  |
| 19 | 1 | 1 |  | 2, 1 |  |
| 27 | 1 | 1 | 3U, 1 | 2 |  |
| 32 | 1 | 1 |  |  |  |
| 33 | 1 | 1 | 4YB | 2 | 1um |
| 39 | 1 | 1 | 3I | 1 |  |
| 40 | 5 | 3 |  |  |  |
| 42 | 1 | 1 | 2HS, 1 |  |  |
| 57 | 2 | 5 | ≥5U |  |  |
| 66 | 5 | 2 |  |  |  |
| 333 | 1x | 1 | 2, 1 | 1 |  |

<sup>a</sup>We characterize an individual as an “immigrant” if it had moved into a study group from elsewhere.
<sup>b</sup>We characterize an individual as being of “uncertain” relationship to breeders if it was caught as a resident independent flier in the territory of one or both of them.

Male post-hatch group auxiliaries included social and genetic sons of both breeders (n = 6 in 2001 and 5 in 2002), an individual home with his father and his replacement mate, an individual home with his mother and her replacement mate, two individuals (one in both years) at home with their half-brothers and their mates, a male that had moved with his older brother, and three immigrants unrelated to both breeders. As such, the male post-hatch group auxiliary population was much more diverse than that of females: 20 individuals (in the two years), all social and/or known genetic daughters of female breeders, of which 18 were also social daughters of male breeders; the two exceptions were a single female (in both years) at home with her mother and a replacement for her father (FX; Appendix A 2b). Within post-hatch groups, genetic relatedness of auxiliaries to female breeders tended to be higher than that for male breeders (Fig. 2).

Post-hatch group size changed in four groups upon the disappearance or death of group members (Table 3): In 2001, the male breeder in one group (of 3) disappeared nine days after hatching had begun in his nest, a 3-year old female auxiliary of another group disappeared 25 days post-hatching, and in a different group, the one-year old female auxiliary and male breeder were killed, in separate incidents, after 15 days of nestling feeding (Appendix A 1a: Group 29). In 2002, a male breeder disappeared after six days of nestling feeding.

## DISCUSSION

The social organization of American Crows in Stillwater, OK, was complex and dynamic, the result of diverse strategies pursued by individuals upon attaining independence, enabled by territory owners across the population. We found little support for classic and current theories regarding the formation, composition, and maintenance of the groups of crows we observed (Conclusions, below), but rather found crows to differ from the majority of other cooperatively-breeding birds in many ways.

Most crows delayed dispersal for at least one year and many remained in natal territories for many subsequent months (one individual in this study was [presumably] still home when at least five years old: PH; Appendix A 2c), yet a few left their natal groups during their first calendar year (one individual in this study dispersed out of his group by the end of November [HE; Appendix A 2e]; his move may have been prompted by human disturbance but others had also dispersed as first-year birds) and a few left in their second year prior to the breeding season. Some individuals that left groups subsequently moved back into them and some moved into other groups (examples in Appendix A 1a and b); some moved in and out of multiple groups (e.g., DE; Appendix A 1a and 2f), some resided with two groups at the same time (e.g., OM and VK, and GN; Appendix A 1b and 2g, respectively), and some did both (e.g., KB; Appendix A 2h). Breeder replacements (of both sexes) resulted in the dispersal of some group members but not all, and the dispersers were not always the same sex as the replacements (Appendix B, Prediction 11). Dispersal of both sexes out of natal and subsequent groups occurred at all times of the year but was most common during the two months or so preceding the onset of nesting across the population, and the week or so preceding hatching within groups. As such, group composition across the population changed most dramatically during January through April of each year, and the compositions of pre-hatch and post-hatch crow groups were highly diverse (Tables 1 and 3).

(On several occasions during late Januarys and early Februarys during years of study [1998-2003], large aggregations of the crows of many groups were observed, in different areas of town; crows at these times were sometimes strangely quiet but were more often making many varied and loud vocalizations and moving around in trees. Although sometimes the eating of late-season pecans took place, foraging was clearly not the focus of crow activity in this context. The function of these seemingly annual aggregations was unknown, and observers speculated that breeding group assignments were being made [and possibly argued about].)

### Intra- and Inter-specific Comparisons

-There are few comparable data available for other populations of crows that live in groups. First- and second-year Western American Crows in Encino, CA, made variable dispersal decisions: about half remained associated with natal areas through their second summers, some left the study area, and some joined the local nonbreeding flock (Caffrey 1992); dispersers dispersed throughout the year, some maintained contact with their social families, and some moved back home after extended periods (Caffrey 1992). Yet during the breeding season, only approximately 37% of pairs had auxiliaries associating with them and 75% of pairs with auxiliaries had only one (and the vast majority of auxiliaries were yearling females [Caffrey 1992]); group size was therefore small, and of all auxiliaries of known dispersal history (n= 28), only one had not hatched in the nest of current breeders (a widowed female breeder who joined another pair and assisted in feeding nestlings). Crows in Encino nested colonially, did not defend territories, and regularly foraged and otherwise interacted in large, loose groups. More similar to our study population in breeding-population social structure, about 80% of American Crow groups in Ithaca, NY, contain auxiliaries (McGowan 2001b, Clark et al. 2006), and group size ranges from 2-12 with a mean of approximately 4 (McGowan 2001b, Townsend et al. 2009). Groups contain auxiliaries of both sexes (equal numbers of yearlings but significantly more male adults than female, and as such, female auxiliaries are significantly younger than male ones [1.45 *vs.* 2.28 years old; Townsend et al. 2009]). Unlike our population, few crow groups in Ithaca contain immigrants (but as in our population, immigrants are unrelated to host group breeders [Townsend et al. 2009]), and aggression among male group members is noted as “obvious” (Townsend et al. 2009); *cf.* the notable *lack* of aggression in our population (Appendix B Prediction 5). Few other details regarding the social behavior of Ithaca crows have been published.

First- and second-year Carrion Crows in northern Spain disperse out of natal groups throughout the year (Baglione et al. 2005) and some dispersers return home after extended periods (Baglione et al. 2006). Approximately 75% of pairs in this cooperatively breeding population (unusual for Carrion Crows) have at least one auxiliary associating with them during the breeding season (Baglione et al. 2006, Canestrari et al. 2008), and group size ranges from 2-9 with a mean of 3 (Baglione et al. 2002a). Auxiliaries include individuals of both sexes that have either delayed dispersal from natal territories (through up to four years; Canestrari et al. 2010) or immigrated into study groups, and the sex ratios of both types of auxiliaries are skewed towards males (Baglione et al 2002b). Approximately half of all groups contain immigrants, all of whom are related to the same-sex breeders of groups into which they move (Canestrari et al. 2008, 2010). Groups are thus extended families (Baglione et al. 2003) wherein relatedness among members, including nestlings, is high (immigrants share in reproduction; Baglione et al. 2002b, 2003). Groups of Carrion Crows are reported to behave as cohesive units, with members that forage close together and rarely leave their territories (especially during breeding seasons; Baglione et al. 2002a, Baglione et al. 2006). Linear food-related dominance hierarchies exist, wherein all males dominate all females (including sons over mothers; Chiarati et al. 2010), and authors report that male breeders “attack” immigrants with greater intensity and more frequently than they do offspring that have delayed dispersal (Chiarati et al. 2011), but also that competitive interactions “rarely escalate into physical aggression” (Chiarati et al. 2012).

Recent information for New Caledonian Crows, which live in small groups but probably do not breed cooperatively (Holzhaider et al. 2011), indicates that offspring may disperse at all times of the year but that at least some remain associated with natal territories for up to “several years” (Rutz et al. 2012). Groups of New Caledonian Crows apparently do not interact much but under certain conditions are known to spend time together (Holzhaider et al. 2011). Mariana Crows (in Rota, Northern Mariana Islands) nest only as single pairs, often renest after failed attempts, and occasionally fledge more than one brood per year (Zarones et al. 2015). One or two young fledge from successful nests (Zarones et al. 2015), and offspring dispersal patterns are variable: some individuals disperse prior to their social parents’ next attempt and the rest are seemingly driven out as breeders begin to nest again, but some first year birds don’t go far and some may return (or visit) after failed subsequent attempts. Small family groups tend to stick together and interactions among groups seem minimal; greater details await further study (S Faegre, pers. comm.).

In contrast, social interactions among crows from different groups in Stillwater were common. In addition to seemingly opportunistic aggregations of members of many groups (in contexts of food exploitation, predator harassment, or other, unknown organizing principles), there were neighboring groups that were friendly with each other and whose members were regularly seen foraging or loafing together, sometimes in the vicinities of each others’ (active) nest trees (e.g., in 2001, Groups 333 and 4, and Groups 10, 42, and 29, and Groups 6, 19, 50, and 45; Fig. 1a). (There were also neighboring groups that mostly ignored each other, and, more rarely, neighboring groups that were NOT friendly and whose members sometimes vocalized at each other and behaved as if agitated [flicking wings and tails while moving among trees], when infrequently in proximity at shared territory boundaries [e.g., Groups 7 and 14 in 2001; Fig. 1].) There were individual crows that regularly visited with “friends” (Silk 2002, Seyfarth and Cheney 2012, Bugnyar 2013) in other groups, and it was not unusual to see unexpected combinations of individuals from different groups just hanging around together somewhere in town.

### Unpaired Crow Decisions

-Early each year in Stillwater, yearling and unpaired adult individuals made decisions about where to live for the next several weeks to months, during the nesting and fledgling stages of breeding seasons. In 2001 and 2002, of 51 pre-hatch auxiliaries with known histories, 71% had chosen to delay dispersal from natal territories (one sixth of them choosing to do so in the presence of a replacement for a parent; four mothers and two fathers), approximately 25% were immigrants, one had moved with a former group member, one had moved with a brother, and one split time between the different groups of his mother and father. For 29% of pre-hatch auxiliaries, including all females not related to female breeders, the criteria upon which their decisions had been made presumably changed as dependent-offspring phases began, because they then chose to change their residency status. Only three times in all the years of our study have we ever seen evidence of breeders possibly disagreeing with unpaired crow dispersal decisions (below [two cases], and Appendix A 2i).

For yearling and unpaired adult crows, options included delaying dispersal (to varying degrees), leaving one group for another, living with more than one group at a time, and temporarily “floating” (in that they appeared to have no fixed area of residence; Koenig et al. 1992, Bruinzeel and van de Pol 2004, Penteriani et al. 2011, Penteriani et al. 2012). (We did not know the destinations of those who dispersed out of our marked population for short or extended periods, yet group living appeared integral to the lives of our crows, and no one we would have described as floating did so for more than a few months at a time.) For auxiliaries, no matter the nature of their relationships with breeders, the potential direct benefits to group membership were myriad.

Although by way of lightening the loads of breeders (Caffrey et al. 2015a) auxiliaries that contributed to nestling feeding may have been delaying their own opportunities to accede to breeding positions (Kokko et al. 2002), many of them opted to do so over the years of our study (yet several did not; Caffrey et al. 2015a). One adult male auxiliary gained ownership of his natal territory (NK; Appendix A 1a), and adult auxiliaries of both sexes -whether related to breeders or not -budded off territories from their groups and bred; some with other group members (e.g., NX and PR [Appendix A 1b], and TF and her mate [Appendix A 2i]), and some not (including cases where within-group potential mates were available, e.g., HE [Appendix A 2e]). Adult immigrant auxiliaries of both sexes replaced missing breeders in their own groups (e.g., KB [Appendix A 2h] and LH [Appendix A 1b and 3c]), and one (not included in this study) replaced the female of a group three territories from hers that had been killed by a Great Horned Owl (*Bubo virginianus*) during incubation. (Presumably, at least some of the previously-unknown individuals that replaced missing breeders in our study groups had also been auxiliaries, in groups outside of our marked population.) During this study, two adult male auxiliaries sired young in their own pre-hatch groups’ nests (NX and KP, in the same group [Appendix A 1b] and likely cases of shared reproduction; Appendix B Predictions 12-15) and two were observed attempting to copulate with incubating females of other groups; one of the latter two (a dispersal delayer) was successful in siring a son, in the nest of a *different* different group (the father of RZ; Appendix A 2e). In two cases in 2001 and 2002, male breeders were assisted in feeding nestlings by male auxiliaries (younger brothers) they had previously helped to raise (NK and EK, and RN and KR; Appendix A 1a, and 2a, respectively), and so past investments were possibly paying off in terms of having their own current nestling-feeding loads lightened. (That RN and KR had moved to a different territory adds a qualifier to a contention of Bergmüller et al. [2007] – that such indirect reciprocity is contingent upon the actor’s attaining breeder status in the same group – and, interestingly, although both EK and KR were unrelated to the female breeders they assisted [each in both 2001 and 2002], their behavior varied considerably with respect to their interactions with their older brothers and their older bothers’ mates [Appendix A 1a and 2j, and 2a, respectively, and Appendix B Predictions 7 and 8, #s 3 and 4].)

Other possible direct benefits to auxiliaries included those associated with site familiarity (Ekman et al. 2004 and references therein) – whether natal or not – and those associated with learning. We saw auxiliaries of both sexes (mostly yearlings) following behind as breeders picked up and discarded sticks during nest building (two were seen to then manipulate and drop a couple of the same sticks) or moved among trees trying out possible sites with first sticks, and sitting near nests and watching as pair members worked on construction or group members fed nestlings. The building of auxiliary nests by auxiliaries (one in particular; Appendix A 2d) – whatever the proximate motivations -also presumably provided opportunities for the learning of breeding skills.

Auxiliaries also benefitted directly in the short term through the food or care provided by breeders; we observed breeders allopreening, feeding, and sharing food with other group members, including immigrants, a few times per month throughout the year (into second years and thereafter) but much more frequently during breeding seasons (auxiliaries sometimes did the same toward breeders, but much more rarely). Notwithstanding food offering may be a “corvid-wide trait” (de Kort et al. 2003), such one-sided behaviors were seemingly not performed in contexts of reciprocity (Stevens and Stephens 2002, De Kort et al. 2006). Food sharing may have been a means of strengthening social bonds (De Kort et al. 2006), but many occurrences were likely adaptive responses by breeders to reduce harassment (Stevens and Stephens 2002, De Kort et al. 2006). Harrassment avoidance has been discussed as a mechanism possibly underlying food sharing (Blurton Jones 1987, Stevens and Stephens 2002, Stevens and Gilby 2004, Hadjichrysanthou and Broom 2012), and our observations supported the hypothesis: crows that had seen another discover something interesting immediately bounded or flew over to investigate. Sometimes discoverers would relinquish partial access without provocation (“latent harassment;” Stevens and Stephens 2002) but sometimes harassers would beg, or (more rarely) act submissively (Verbeek and Caffrey 2002), before being allowed access or fed. Some bouts of allopreening by breeders were also likely means to reduce harassment: auxiliaries sometimes solicited the behavior and were sometimes rewarded. Occasionally, though, if breeders declined to allopreen and began to move away from solicitors’ bent-over bodies, solicitors used their feet to pinch the toes of breeders; this sometimes got breeders to change their minds, but other times got solicitors pecked on their heads before breeders flew off. (In a similar context in Encino CA, a solicitor stepped on the foot of the breeder as the latter attempted to move away, pinning her in place. This solicitor received a peck on the head and no allopreening [CC, pers. obs.].) At other times, though, mostly at nests and by female breeders to auxiliary members of post-hatch groups who fed nestlings, allopreening bouts were unsolicited, long, and gentle; different in nature than ostensible harassment-reduction responses (and likely not a mechanism of recruitment, since recipients were already feeding nestlings), possibly a means of encouragement or reward (Bergmüller et al. 2007), and seemingly, to us, pleasurable to both parties. (As allopreening is often directed to recipients’ heads [where they themselves cannot reach], it likely provides advantages by way of reducing parasite loads [De Kort et al. 2006]. Allopreening may also provide physiological benefits, similar to those resulting from allogrooming in nonhuman primates [Aureli and Schaffner 2002 and references therein], including increased endogenous brain opioid release likely resulting in “pleasant sensations” [Aureli and Schaffner et al. 2002].)

With regard to potential *indirect* benefits, auxiliaries did not, on average, enhance the production of nondescendant kin (at least in terms of the number of young fledged per nest) via their presence or feeding contributions in 2001 or 2002 (Caffrey et al. 2015a), yet for those whose parents chose to compensate with their assistance (Caffrey et al. 2015a), such benefits would theoretically accrue in the future.

Fewer ideas are available regarding possible benefits underlying the decision by some auxiliaries to disperse out of groups at hatching. The obvious explanation -so as not to have to pay the costs associated with feeding nestlings – did not seem sufficient: the highly variable patterns of nestling feeding by auxiliaries in this population included some in which zero to few trips were made during many hours of observation (Caffrey et al. 2015a), and there was no evidence of false feeding (expected if auxiliaries wanted to be perceived as meeting some minimum level of investment; Young et al. 2013) or of punishment or retribution (Bergmüller et al. 2007, Cant 2011, Young et al. 2013) by breeders toward auxiliaries contributing little to nothing toward nestling care (Caffrey et al. 2015a). Thus dispersers did not appear to be leaving to avoid the high “performance costs” (Barclay and Reeve 2012) of staying (and, in fact, appeared to be leaving their “ideal” worlds [Bergmüller et al. 2007]; being able to use the resources of territories but not having to provide “help”). Yet given the potential advantages to establishing social relationships and gaining familiarity with potential breeding areas (below), it is possible that dispersers left to avoid the “opportunity costs” (Barclay and Reeve 2012) of staying: nine of seventeen marked dispersers did *not* return to their groups, including two individuals known to be alive weeks after dispersal and three others after more than a year (Appendix A 3); presumably some of their floater experiences were successful and they ended up settling, and possibly breeding, elsewhere (*sensu* Penteriania and Delgado 2012). For females, post-hatch residency decisions were clearly tied to indirect benefits: all females unrelated to female breeders dispersed out of groups for nestling (and fledgling) periods, and all of those who stayed were genetic and social offspring of at least female breeders (and for all but one, male breeders as well). In addition to the possibility that dispersers were avoiding increasing the fitness of unrelated females (again, breeders tended to reduce nestling feeding contributions when assisted by auxiliaries; Caffrey et al. 2015a), this observation aligns with decisions made by female Seychelles Warblers to only help at nests attended by their social mothers (Richardson et al. 2003, Komdeur et al. 2004), in support of the contention of Heinsohn and Legge (1999) and Heinsohn (2004) -based on Cockburn’s (1998) finding of a potential sex-biased effect of auxiliaries on productivity -that female helpers primarily seek increased indirect benefits via kin selection (when helpers are predominantly female or are of both sexes, nest productivity is increased; Cockburn 1998), whereas male helpers, whose contributions are usually not associated with increased fledging success (Cockburn 1998), instead seek opportunities for direct reproductive benefits. Yet that some female crows chose to leave natal groups with mothers present, and thereby pass on opportunities for increased indirect fitness, indicates that not all them were marching to the same selective drummer.

### Breeder Decisions

-For breeders, there were tangible benefits associated with having extra individuals assist in nestling care (breeders tended to compensate for auxiliary contributions; Caffrey et al. 2015a). Breeders also stood to gain by way of the potential benefits associated with offspring delaying dispersal for one or more years, including improved nutritional condition for *themselves* during winter months, as a possible result of increased foraging efficiency or predator detection (Pravosudova and Grubb 2000). Potential benefits would also include increases in survivorship of offspring and/or in their chances of securing successful breeding opportunities in the future (Ekman et al. 2004, Pruett-Jones 2004, and references therein), and any benefits associated with their direct involvement in nesting attempts (e.g., benefits involving the learning of nestling-care details [Komdeur 1996, Cockburn 1998, Dickinson and Hatchwell 2004]). We saw evidence of one possible route to increased survivorship for offspring who chose to remain (or to return to natal territories after having previously dispersed): during some winters, when particularly heavy storms left the ground covered with snow for two or more consecutive days, crows shifted more of their foraging attention to tree canopies, and first- and second-year auxiliaries tended to follow after foraging adults under such conditions. On several occasions, observers saw breeders retrieve (apparently) cached pecans and other food items from branch crotches covered in snow (crows had been “traveling mentally in time and space” [Emery and Clayton 2004]) in front of other group members (and on occasion share such items with them; a means of increasing offspring survival in Siberian Jays, *Perisoreus infaustus*; Ekman and Rosander 1992). We also observed a father and sister (a brood-mate) spending time with and feeding an injured female in February of her second year (before she disappeared; Appendix A 4).

Less clear were the potential benefits to breeders of allowing unrelated individuals to move in – individuals old enough to threaten breeder fitness and who would likely move out for periods of nestling and fledgling care. Only once between fall 1997 and December 2002 did we observe an incipient immigrant not being accepted readily by breeders: an adult female moving in during the month including nest completion, egg fertilization and laying, and the onset of incubation (DE as she moved into Group 19 in March of 2000; Appendix A 1a and 2f). In another case, an immigrant of several months (LH; an adult female) was observed being chased by breeders, shortly after incubation had begun (seemingly in the context of a conflict with neighbors [Group 27 in 2002; Appendix A 1b]). The otherwise easy acceptance of immigrants and visitors by breeders and other group members, and the sharing of food with immigrants by breeders (above), suggested that any costs associated with competition for resources were easily overcome by benefits associated with allowing extra-group individuals to move in for varying amounts of time (group members should prevent such behavior if they suffer competition costs; Griesser et al. 2008), but what might have been the possible “returns” on breeder “investments” (Bergmüller et al. 2007) in such individuals?

Across the population, group size did not appear to influence a pair’s ability to obtain or maintain ownership of breeding space; the budding of territories by new pairs happened regularly, and pairs without auxiliaries were not aggressed or encroached upon, or otherwise threatened by other population members. Canestrari et al. (2007, 2008) suggest that larger Carrion Crow groups may be better able to thwart predation as a possible explanation for the acceptance of individuals that do not contribute to nestling feeding (mostly “subordinate” females [Canestrari et al. 2008]), but the authors speak only to breeding seasons, and because Carrion Crow immigrants are related to breeders of the same sex and share in parentage of nestlings (Baglione et al. 2002b), the interests of Carrion Crow breeders differ considerably from those of the breeders in our population. Still, the benefits of larger groups in contexts of predation are widespread, many being realized through reduced vigilance rates (Proctor et al. 2006, Beauchamp 2008) which may then translate into increased foraging rates (Proctor et al 2006, Fernandez-Juricic et al. 2007), and crows in larger groups in Ann Arbor, MI, spent less time vigilant (scanning) and more time foraging than those in smaller groups (Ward and Low 1997). Predation did not cause high mortality in our population, and most of the predation that did occur (of which we were aware) was by Great Horned Owls -nocturnal predators unlikely to be influenced by crow group sizes – however, that there are potential advantages to spending less time vigilant cannot be denied.

Larger groups are also associated with increased foraging success for many bird species (Beauchamp 1998). Crow group size did not affect a group’s ability to defend against the occasional intrusion into territories of large numbers of crows -many of which were likely non-resident migrants -when pecan trees were fruiting [October-December]), nor did it affect an individual’s ability to exploit opportunistic bonanzas of food that became available outside of territories in Stillwater (e.g., dumped agricultural refuse). Crows did most of their foraging, however, at home and in the company of group members -just how they lived most of their diurnal lives, throughout the year, no matter the weather or foraging conditions. Possibly, it was the nature of the crow groups themselves, and the dynamics among them, that enabled this cooperative breeder to thrive in the northern hemisphere, where the harsh winters are thought to render allowing individuals to share territories disadvantageous to breeders (Ekman and Rosander 1992, Ekman et al. 2002, 2004 and references therein). The large brains of crows (Emery and Clayton 2004, Emery et al. 2007) are associated with the ability to respond adaptively to novel situations and environmental variation (Lefebvre et al. 1997, Sol et al. 2002, 2005, 2007), and the pace at which change can occur in the human-modified habitats in which they live can be rapid (Wong and Candolin 2015). We suggest the crows in our population may truly have been benefitting from group augmentation (Wright 2007): the fitness of individual group members was increased by way of their relationships with other group members.

There were numerous potential advantages to breeders (and their offspring) to including in their groups individuals that varied in experiences, skills, personality or temperament traits (Reale et al. 2007, Scheid and Noe 2010, Ensminger and Westneat 2012), foraging tactics (Bolnick et al. 2003, Rockwell et al. 2012), and innovativeness (Lefebvre 1997, Reader 2003, Bókony et al. 2014), including increasing the groups’ efficiency at problem solving (the “pool-of-competence effect,” Morand-Ferron and Quinn 2011; Griffin and Guez 2015 and references therein): some immigrants may have been particularly capable of locating new food sources (“producers;” Arbilly et al. 2014 and references therein) or of devising new ways to exploit resources already available (Liker and Bókony 2009). Individual crows in groups with varied others likely experienced reduced neophobia (Katzir 1982, Stöwe et al. 2006, Chiarati et al. 2012 and references therein) and/or increased object exploration (Katzir 1982, Miller et al. 2014); it may even have been the case that including “initiators” in their groups allowed breeders to be more conservative in risk taking (Katzir 1982). Group members may also have benefitted via increased success through social facilitation in contexts of high-risk prey species (Miller et al. 2014), and possibly also reduced predation risk, through the innovativeness of individuals (adaptive also in the context of responses to enemies; Sol et al. 2007) and the use of “public” information (Danchin et al. 2004). One strategy to cope with environmental challenges successfully employed by cognitively-capable animals (including New Caledonian Crows [Holzhaider et al. 2010] and Common Ravens [Fritz and Kotrshal 1999, Schwab et al. 2008]) is social learning (Coussi-Korbel and Fragaszy 1995, Alvard 2003, Laland 2004); through copying and “observational learning” (understanding the intent or goal of the demonstrator; Alvard 2003), individuals can add options to those available through personal experience and then selectively choose among them under different conditions so as to maximize outcomes (Laland 2004, Kendal et al. 2005, Grüter and Leadbeater 2014). (Breeders may also have benefitted by reducing costs associated with defense of the resources they chose to share [Stevens and Stephens 2002 and references therein] in winters; during springs, possibly, enough food was available such that defense was not necessary [*sensu* Penteriani and Delgado 2012]).

Knowing others through shared group membership also sets up relationships with value (Roberts 2005, Bergmüller et al. 2007, Fraser and Bugnyar 2010a), relationships that might somehow provide advantages in the future (a kind of “deferred intraspecific mutualism” [Clutton-Brock 2002], or “inter-dependence” [Roberts 2005]). Such relationships, including friendships, may translate into measurable benefits such as access to food (Braun and Bugnyar 2012), increased survivorship (Silk et al. 2010Barocas et al. 20011), or increased reproductive success (Silk 2002 and references therein, Silk et al. 2003, 2009, Cameron et al. 2009, Schulke et al. 2010, Massen and Koski 2014 and references therein). There are also intrinsic benefits to be gained through socializing, and through cooperating (Schuster and Perelberg 2004, Bergmüller et al. 2007), benefits possibly mediated proximately through something akin to “pleasure” (Aureli and Schaffner 2002, Schuster and Perelberg 2004) but with eventual reproductive payoffs (Schuster and Perelberg 2004). We saw the manifestation of two types of ways in which getting to know immigrant individuals was advantageous: breeders of host groups ended up in long-term relationships with both female and male immigrants through the latter’s becoming either replacement mates (e.g., LH and KB in this study; Appendix A 1b and 3c, and 2h, respectively) or neighbors (via budding, e.g., PR and HE in this study; Appendix A 1b, and 2e, respectively). In Red-winged Blackbirds (*Agelaius phoeniceus*), familiarity with neighbors has been shown to be associated with enhanced reproductive success (Beletsky and Orians 1989).

There are immediate health-related advantages to friendships, as well (Uchino et al. 1996, Aureli and Schaffner 2002, Silk 2002 and references therein), including reduction in the day-to-day stress of the friends involved (Wittig et al. 2008, Crockford et al. 2008, Bugnyar 2013), stress potentially capable of negatively influencing, among other things, problem-solving performance (Bókony et al. 2014).

And so there exist ideas and examples of possible advantages to breeders for the seemingly unusual decisions they made regarding the availability of their groups to others, and because auxiliaries only rarely succeeded in parenting fledged offspring (only a single case in 2001 and 2002 [above, the father of RZ; Appendix A 2e]; we suspect the extra-pair young in Nest 333 in 2001 and 2002 [above, NX and KP; Appendix A 1b] were sired through shared reproduction [Appendix B Predictions 12-15]), there do not appear to have been many reproductive disadvantages to allowing immigrants to move in or to allowing unrelated step-offspring to remain.

### Conclusions

-Among the networks of genetic and social relationships in our study population, we did not find much support for extant theories of group formation, composition, and maintenance in cooperatively-breeding birds. Rather, our study population differed considerably from those described for most other cooperative breeders, including other crows. Though ostensibly classically territorial, our crow groups exhibited relaxed territoriality and the members of many were friendly with members of others. Inter-group movement of unpaired individuals was common and mostly uncontested, and intra-group aggression was rare (Appendix B Prediction 5). Widowed female breeders were not evicted (Appendix B, Predictions 7 and 8, and *cf.* Emlen 1995, Cockburn 1998) and the moving in of replacement breeders did not always trigger dispersal out of groups by auxiliaries of the same sex (Appendix B Prediction 11, and *cf.* Emlen 1995, Ekman et al. 2004, Koening and Haydock 2004). Members of the diverse pre-hatch auxiliary population did not appear to be making the best of a bad job (Koenig et al. 1992, Emlen 1994, Hatchwell and Komdeur 2000, Bergmüller et al 2007, Lenda et al. 2012 and references therein), or to be basing their dispersal decisions on such factors as “a special value to home” (Ekman and Griesser 2002 and references therein, Ekman et al. 2002, 2004, 2006, Chiarati et al. 2011), the presence of nepotistic parents (Brown and Brown 1984, Ekman et al. 2001, Chiarati et al. 2012, Ekman et al. 2004 and references therein), or even *any* parents (*cf.* Ekman and Griesser 2002). That immigrants left (their own) tolerant parents at home contradicts the prediction of Covas and Griesser (2007) that given delayed breeding, if parents are tolerant, delaying dispersal by offspring should be the “best strategy.” Prior to 2003 (before the first episode of high West-Nile-virus-related mortality in the fall of 2002; Caffrey et al. 2005), few auxiliaries became breeders in their own groups, and so independent crows were likely not choosing where to live on the basis of queue length (*cf.*, e.g., Wiley and Rabenold 1984, Williams and Rabenold 2005, Bergmüller et al. 2007), and the territory-quality considerations that dominate the theoretical literature (e.g., Koening et al 1992, Ekman et al. 2004 and references therein) were also unlikely to have been important determinants of the decisions of unpaired crows in our population: crows moved freely into and out of our population, space was available each year (into some of which crows eventually moved during our study), budding was common, successful pairs did not require a lot of space (e.g., Group 32 in Fig. 1), population members were able to exploit temporarily abundant resources (e.g., pecans) in territories other than their own, and pre-hatch auxiliaries did not appear to be choosing groups based on relatedness to breeders or breeders’ reproductive success (often equated with territory quality, e.g., Koening et al. 1992, Kokko and Lundberg 2001). The imminence of hatching and nestling-feeding stages of breeder reproductive attempts apparently changed conditions enough for some auxiliaries that higher benefits were to be gained elsewhere and they moved out, at least temporarily. This round of decisions *was* influenced by genetic relatedness, at least for females, who uniformly dispersed out of pre-hatch groups if unrelated to female breeders. Eight of seventeen marked dispersers returned (Appendix A 3), presumably not having found better options elsewhere.

Given the long lives of crows (in captivity, up to almost 60 years [Associated Press 2006]; in the wild, up to into their twenties [K. McGowan, pers. comm.]), their pre-breeding dispersal and residency decisions from year- to-year were no doubt being made in long-term contexts. Given their delayed maturation and the benefits to doing so (Covas and Griesser 2007 and references therein, Penteriani et al. 2011 and references therein), most crows put off pairing and establishing territories for a few years after maturation (some did so at three- and four-years old, but others were still unpaired at four and five). During this time – while at the same time forestalling the risks (to future reproductive potential) associated with current breeding – crows were presumably preparing to exploit worthwhile breeding opportunities in the future, through adding knowledge (including that underlying “informed dispersal;” Clobert et al. 2009), skills, networks of social relationships, and savvy to their “toolboxes,” through living, interacting, and cooperating with others, to some extent regardless of genetic relationships (rare among vertebrates; Clutton-Brock 2009).

Clearly there were advantages to future breeders, particularly as opportunistic generalists, to moving around their population and getting to know the resources and dangers associated with potential breeding areas. It’s pretty clear, too, there were advantages to moving around their population and getting to know other crows. Investment in social relationships allows interactants to predict the actions and responses of partners, making interactions more efficient (Aureli and Shaffner 2002, Alvard 2003) and reducing stress (Crockford et al. 2008). As such, territory owners (and their relatives) stood to gain as well, by establishing relationships and making friends with unrelated group members, so as to be able to maximize the benefits and minimize the losses associated with current and future relationships (e.g., in contexts of foraging, cooperative defense, mate choice, and budding and subsequent neighbor status). There may have been benefits, too, to crow sociality itself; in feral horses (*Equus caballus*), e.g., juveniles with more associates were more likely to survive a catastrophic event (Nuñez et al. 2014). If so, then it may have been the case, as Nuñez et al. (2014) suggest for horses, that the more individually varied the group, the more advantageous regarding the social development of young crows. As such, this would provide an additional route to benefits to breeders welcoming others into their groups.

It thus appears that the shared interests in high-quality relationships (Silk 2007, Fraser and Bugnyar 2010a) among individuals of different sex and status in this population likely contributed to the surprisingly fluid and benign nature of its society.

### Emlen’s Predictions

-Drawing from kin selection, ecological constraints, and reproductive skew theories, Emlen (1995) made cases for 15 predictions having to do with the formation, organization structure, stability, and social dynamics of families, defined as groups created when grown offspring continue to interact regularly with parents. The logic of his predictions was based on imagining groups of breeders and their genetic relatives (almost exclusively; only replacement mates were considered as unrelated group members), yet the applications of his logic can be extended, to some extent, to the groups of crows of varied relationships in Stillwater; to wit, group formation occurs when the long-term interests of unpaired individuals is best served by living in space owned by others. Some of the data herein, including some in Appendix A, lend themselves to placement in the contexts of predictions # 2, 5, 7-9, and 11-15 of Emlen (1995); those related to group formation, group dynamics associated with breeder deaths and replacements, within-group aggression, and the sharing of reproduction within groups. We include this treatment as Appendix B.

## ACKNOWLEDGMENTS

We are extremely grateful to A. K. Townsend, L. M. Stenzler, and I. J. Lovette for genetic work, and we thank R. Van Den Bussche, R. Kimball, and J. Dickinson for assistance with crow sex determination and DNA isolation and storage.

Many hundreds of volunteer hours went into getting crows marked, their sexes determined, and their DNA isolated, in addition to those that went into collecting the data reported here: thank you to every one of the many family and friends, and the friends and family of family and friends, who assisted us. We are especially thankful for the extraordinary selflessness of D. M. Woods and S. Fierer, and grateful for field-related help from S. C. R. Smith, P. O’Malley, H. Moravec, M. Disney, M. Steele, K. Adams, R. Soto, J. and N. Wilhm, T. Pappan Turnipseed, K. O’Malley, J. Moravec, and C. Lampe, and, in memorium, P. Underhill and M. Hryzrn. We are also especially thankful for T. W. Hackler’s assistance in the field and with sex determination and DNA isolation.

For assistance with facilitating the capture of crows and getting to crow nests, we thank the Arbortec Tree Expert Company, the City of Stillwater Parks and Recreation Department, the City of Stillwater Police Department, the City Manager’s Office of Stillwater, Oklahoma State University (OSU) Physical Plant, OSU Police Department, and OSU President and Mrs. J. E. Halligan. We also sincerely thank the many Stillwater residents who let us set up capturing devices in their yards, pull cherry pickers onto their lawns, and climb and trim their trees, and who warmly invited us into their homes and lives.

We gratefully thank K. McGowan, A. Clark, and W. Koenig for stimulating conversations over many years and assistance of all kinds, S. Faegre for information on New Caledonian and Mariana Crows, S. Beaudoin for information on the dispersal of one of our crows to Pawnee, OK, M. Wehtje for Figure 1, and several reviewers of past drafts of this work for their helpful comments.

I (CC) wish to also acknowledge the members of our study population; individuals who earned the deep respect of my students and me, who exasperated us, entertained us, and made us think a lot about what and how they might be thinking. Every one of them ended up either dying the grueling, painful death of West Nile virus infection themselves, or watched, and mourned, as most of their family members and friends inexplicably did so. Witnessing their suffering had been heart-breaking; imagining it unbearable.

This work was funded in part by grants from the Payne County Audubon Society, and gen-erous assistance from J. and N. Wilhm, and E. and H. Caffrey.

## APPENDIX A.

### Some details

#### 1. Relationships within and among crow groups

a. Involving Groups 111 (extinct), and 10, 11, 19, 29, 42, and 92 (Fig. 1). The players:

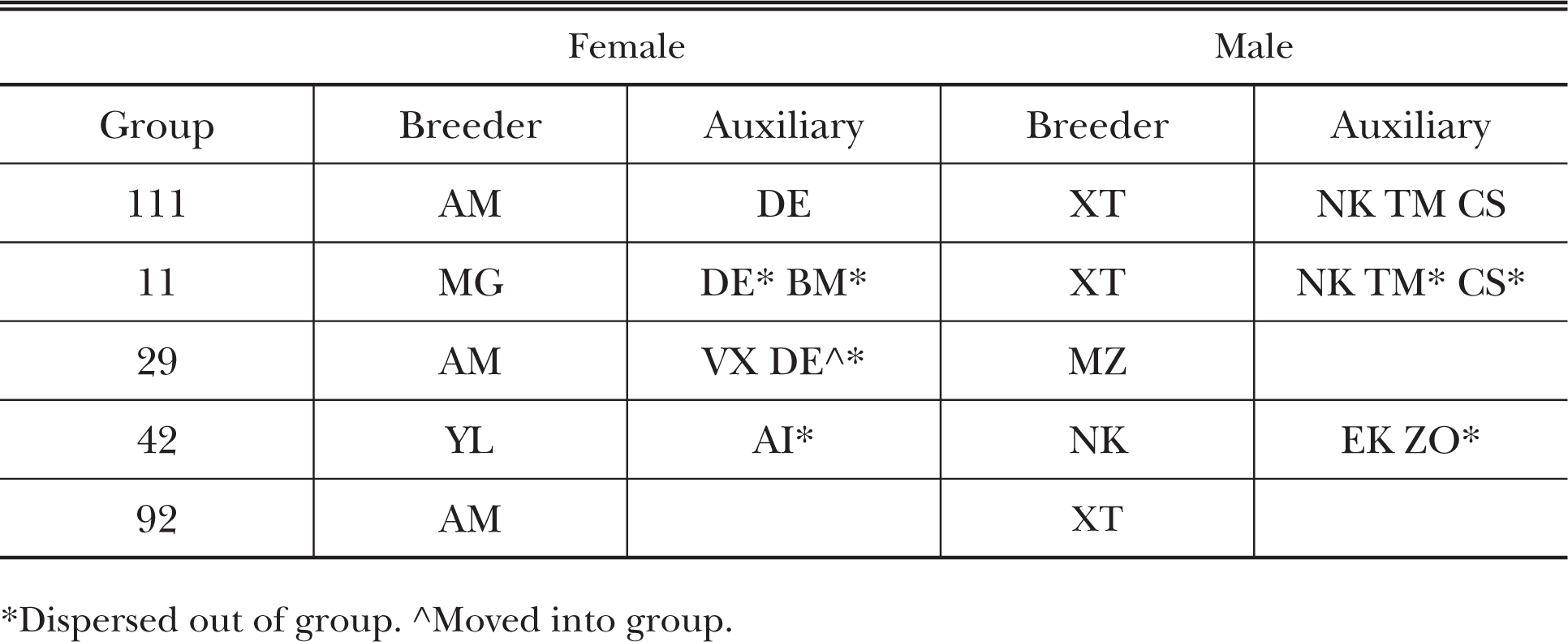 In early March 1999, an injury befell the female (AM) of Pair 111 (a group no longer extant by 2001 but that lived in the same general area occupied by Group 42 in Figure 1a; AM had been shot with a pellet gun) and she was removed from the field for several weeks. Within a few days, a replacement female (MG) moved in and paired with AM’s mate (XT, creating Group 11 [Appendix C 1ii]); all four of AM and XT’s one-year old offspring (a female -DE - and three males: CS, TM, and NK) remained home through MG and XT’s first attempt (abandoned 5-6 days after hatching) and the building of their second nest. At around the beginning of incubation of the 2^nd^ attempt, TM (below, 2d) moved out and into neighboring Group 10 (Fig. 1 Appendix C1iii; he remained with Group 10 through 2002; r with both breeders = 0). DE, CS, and NK remained and fed the nestlings of MG and XT, as did TM the nestlings of Group 10. We were aware that AM had returned to the general area upon her release but could not accurately track her whereabouts until November, when DE left Group 11 and joined AM (her mother) and her new mate (MZ; r with XT = 0.5207 [Appendix C1iv]), with whom DE remained until March 2000. DE then moved in with Group 19, with whom she resided until February 2001. AM and MZ (Group 29) nested next-territory to MG and XT in 2000 (NK, and MG and XT’s yearling female daughter [BM] were observed feeding nestlings; CS, still at home, was not [Appendix C1v]). In June of that year MG disappeared (Appendix C1vi). CS and BM dispersed out of Group 11 in September (Appendix C1vii). In October, an adult female (YL) moved in (Appendix C1viii). As the breeding season of 2001 approached, XT and NK (father and son) seemingly amicably vied for access to YL (Appendix C1ix), and both were observed sitting and allopreening with her (once with her between the two of them); she seemed to us to prefer NK. One day XT disappeared (two days after we had caught and handled him) and by the next day NK and YL (Group 42 [Appendix C1x]) were working on their nest. NK’s two yearling half-brothers (EK and ZO) stayed and helped raise his nestlings; his yearling half-sister (AI) moved out at hatching (and moved to Pawnee, OK; 44.9 km away [below, 3f3, Discussion, and Appendix C1xi]). Next territory, Group 29 consisted of AM, MZ, and their yearling daughter (VX), and was being visited regularly by a 2-yr old female auxiliary from Group 10 [Appendix C1x]. When the nestlings of Group 29 were about 2.5 weeks old, VX was killed by a Great Horned Owl (*Bubo virginianus*) outside of town. A day or two later, MZ was hit and critically injured by a car; he remained alive but unable to get to his nest, and impossible to catch, for almost two weeks. Within a day of MZ’s injury, XT was observed attempting to copulate with AM at her nest (presumably, she was *not* fertile). AM fed nestlings on her own for more than a week before we provided her with food. Two days after MZ was caught and removed from the field, XT moved in with AM (it had been six weeks after his departure from his and AM’s original territory, now owned by their son [NK; Appendix C1xi]). During two final watches at AM’s nest, XT was observed to feed the single remaining nestling several times. (All four of AM and MZ’s nestlings were fathered by MZ.) AM and XT remained paired and bred together again (Group 92) in 2002, next territory to two of their sons: NK (and YL and their yearling son, and EK [below, 2j]; ZO had moved out [Group 42]) and TM (still in Group 10 [Appendix C1xiv]). (NK was observed to land on the rim of Nest 92, to look in briefly [while his parents were temporarily away] on the day first day of hatching.)
b. Involving Groups 3, 333, 4, 14, 27, and 55 (Fig. 1). The players:

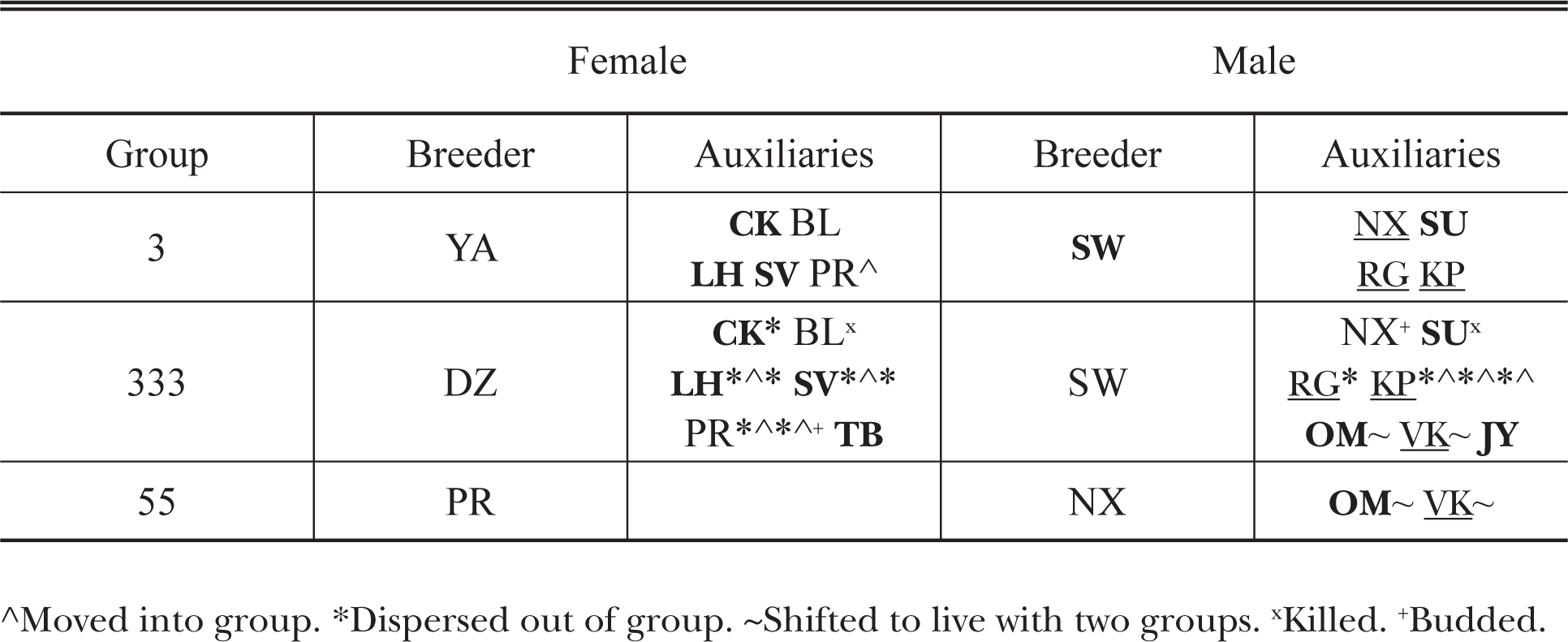

##### SW is father. <u>NX</u> is father

During the nesting season of 1999, Group 3 (a group no longer extant by 2001 but which lived in the same general area occupied by Group 333 in Fig. 1a) consisted of two breeders (YA and SW) and four auxiliaries (an adult male, NX [*not* the genetic son of either breeder: r with YA = 0.0698, with SW = 0.2932], and three one-year olds caught the previous fall; two females (CK, BL) and a male (SU), two of which were the offspring of YA and SW [we had no genetic data for BL; Appendix C2ii]), all four of which fed nestlings. Four young fledged, survived, and remained home thereafter: two females (LH and SV, both daughters of YA and SW) and two males (KP and RG, both sons of YA and NX [the auxiliary; Appendix C2iii]).

In January 2000, a third-year female (PR) from Group 14 (one territory away; Fig. 1a) moved in with Group 3; group size was then 11 (Appendix C2iv). In March, YA disappeared; an adult female (DZ) moved in - and although it seemed clear DZ was pairing with SW (creating Group 333), NX was often observed in their proximity in the early days of her residence - and LH, SV, and PR moved out (Appendix C2v). Six auxiliaries remained through fledging (their nestling-associated behaviors could not be determined because of poor viewing conditions), after which CK moved out. Three young fledged, two males (OM [the son of DZ and SW] and VK [the son of DZ and NX]) and one of unknown sex and parentage (we were unable to catch this individual). In October, LH and SV moved back in, and KP and RG moved out to join Group 4 (one of Group 333’s next-territory neighbors [Fig. 1], and a group with which Groups 3 and 333 were friendly; r with both breeders essentially zero for both KP and RG [Appendix C2vi]). In November PR moved back into Group 333 (Appendix C2vii). RG remained with Group 4 through hatching in 2002 (1.5 years later).

KP moved back into Group 333 over January-February of 2001; group size was then 12, briefly, before the yearling of unknown sex and parentage disappeared (Appendix C2viii). All nine other auxiliaries remained until the week or so preceding hatching in Nest 333, at which point LH, SV, KP, and PR left the group (Appendix C2ix; all were regularly seen thereafter in surrounding areas, i.e., they appeared to be “floating,” in the sense of having no fixed area of residence [Koenig et al. 1992, Bruinzeel and van de Pol 2004, Penteriani et al. 2011, Penteriani and Delgado 2012]). Three of the five remaining auxiliaries fed nestlings: two males (NX and OM), and BL. Three young fledged (one having been sired by KP, who had moved out at hatching), but one (KP’s daughter) died shortly thereafter (Appendix C2x). By the fall of 2001, PR and KP had moved back into Group 333. LH was moving in with Group 27, several territories away (Fig.1a). SV (below, 3f2), no longer seen in areas around town, visited Group 333 infrequently but regularly (2-3 times/year). SU was killed by a predator – we found his banded leg – sometime in December.

Early in 2002, PR and NX were beginning to pair (Appendix C2xii). KP, OM, and VK were all observed also attempting to spend time near PR. By late February, PR and NX (Group 55) budded off a territory from that of Group 333 (Fig. 1b), and OM moved with them (VK split his time between Groups 333 and 55 -those of his mother and father (Appendix C2xiii), although his “home” group was 333 [that of his mother and her mate]). PR and NX abandoned a first nest and began working on another, a period during which OM was observed working on a separate nest. PR was seen to make a trip to OM’s nest during one of our nest watches (she carried to it and worked in a stick); during the same watch (= 150 min) NX made eight trips, all bringing nothing and four of which involved his removing material (three sticks and a string) and carrying it away. Seven of NX’s eight trips to the nest were made when OM was present and OM always left at NX’s arrival; only twice did he stay nearby and return to adjust material after NX’s departure. OM abandoned his nest and infrequently succeeded at helping PR and NX build theirs; if anywhere around when OM arrived at the nest, NX would immediately approach and chase him away. OM did not seem particularly interested in being close to PR, and his interactions with NX away from the nest appeared normal. Once incubation was underway, OM and VK visited the nest occasionally, uncontested by NX. PR and NX abandoned this attempt after 18 days of incubation.

Next territory, Group 333 included the breeders and four auxiliaries: KP, VK, and two one-year olds (a male and a female [Appendix C2xiii]). KP moved out at hatching (Appendix C2xiv) and, with an unmarked crow, built a nest about 150m away from Nest 333. The male breeder of 333 (SW) disappeared after about a week of nestling feeding at his nest, and a week later KP was observed feeding nestlings there; this was two days before incubation began at his own nest, which was abandoned by a week thereafter. KP then moved back into 333 [Appendix C2xv]. Although never subsequently observed feeding nestlings or fledglings (of which there were two surviving ones, both sired by SW), KP sat with them and peacefully tolerated their begging. OM moved partly back in with Group 333 after Nest 55b was abandoned (his “home” group remained 55; that of two former group members to whom he was not related); he and VK were then both members of the two Groups 333 and 55 (Appendix C2xv), yet whereas OM was never seen at Nest 333, VK made a large proportion of nestling feeding trips subsequent to the disappearance of SW (in direct contrast to his behavior the year before: although often observed with group members away from the nest, VK was observed to make only one brief visit to the rim during the nestling period [Caffrey et al. 2015a]).

Across town (Fig. 1b and Appendix C2i), Group 27 included five auxiliaries, including LH (who had moved in the previous fall) and a two-year old female (DX, the daughter of the female breeder; the current male was likely a replacement for her father) who had (apparently) moved out as nestlings fledged the year before and was not seen again until February 2002, when observed a few times in large aggregations of crows outside of town. (As she moved in, in March, on two occasions we saw the male breeder and her half-brother move toward her in ways that caused her to move back a bit, but their lunges and hops seemed lackadaisical and DX did not behave submissively, and moved only temporarily, before everyone resumed foraging.) LH was observed several times foraging and loafing with members of Group 27, including breeders, as they built their nest and began incubation, and a few times alone with an adult male auxiliary in the group; they spent time in an area bordering Group 92 (Fig. 1b and Appendix C2i) and the male was once seen prospecting potential nest sites with a stick. On three occasions we observed territorial disputes in this area; LH and the male auxiliary were chased out by Pair 29 once, and twice involved all six adults: the first resulting in Pair 29’s retreat and Pair 27 turning to LH and chasing her from the area, too, and the second, a week later (after the female breeder of 27 [HO] had already begun incubation, *not* nearby), when all six dispersed from the area. LH then moved out. HO was one of the first of approximately one-third of our population to die that fall in the wake of West-Nile-virus’ initial arrival (Caffrey et al. 2005), at which point LH moved back into Group 27, paired and ultimately (in 2003) bred with the widowed male.

#### 2. Individuals

a. RN: RN lived with his parents, in Group 1 (a group no longer extant by 2001 but which lived in the same general area occupied by Group 41 in 2001 [Fig. 1a]; it varied in pre-hatch size from 4-10 in the years 1997-2000), through the end of his fourth year (1999). Group 1 was friendly with their neighboring group to the northwest (Group 16; no longer extant by 2001 but which lived in the same general area occupied by Group 33 in 2001 [Fig. 1a]); members of both groups were regularly observed foraging and loafing in close proximity. Near the end of 1999, RN and a younger brother in his second year (KR) were observed a few times in small, casual groups of members of Groups 1 and 16 and 19 (Fig. 1a), including the female breeder from 16 but not the male. RN moved in with the (presumably widowed) female breeder (GP, creating Group 33) in early 2000, and KR moved with him. We never saw *any* paternity-protecting or -seeking behavior by RN or KR during the nesting season of 2000 (or 2001 or 2002), and KR assisted in feeding nestlings and fledglings, of which there were three. One of the three was shot and killed in the backyard of a Stillwater resident in December 2000, and one disappeared during the following February (a popular dispersal time). RN, GP, KR, and one yearling all fed the nestlings in Nest 33 in 2001; a nest physically damaged by an unknown process late in the nestling stage (likely storm- or predation-related) and from which one young fledged. This group of five was particularly close-knit and often seen together, and together they fledged five young in 2002. By the end of that September, of this extended family of 10, only RN and two young of the year had survived the West-Nile-virus epidemic [Caffrey et al. 2003]. The two second-year individuals dispersed in early 2003 as RN paired with KV, a four-year old female that had remained in her natal territory (next to that of Group 33: Group 19, a host group to many visiting crows over the years [both short- and long-term] and whose members were friends with members of Groups 1, 16, and 6) and assisted in raising nestlings each year. RN and KV fledged four young in 2003 and by the end of the summer, all six members of this family had disappeared; likely the victims of the second, devastating round of West-Nile-virus-related mortality (approximately 65% of our population died during the late summer and fall of that year [Caffrey et al. 2005]).
b. FX: FX was one year old and living with both parents when KB (an adult male; below, 2h) moved into her group (#15, a group no longer extant by 2001 but that had lived in the same general area occupied by Group 115 in 2001 [Fig. 1a]), in January 2000. Her father, suffering from a leg affliction, disappeared by the end of the year and KB paired with FX’s mother (JN; Group 115) in 2001. FX remained connected to her natal territory but left for several weeks in the fall of 2000, when she was observed in the territory of a group (#1; above, 2a) in which the female breeder was suffering greatly from a leg injury. We caught FX with members of Group 1, which (the capture) may have been the reason she then moved back home. (The female breeder of Group 1 died shortly thereafter and a second-year [sexually immature] female auxiliary from Group 14 [Fig. 1] moved in with the widowed male.) FX had assisted in feeding nestlings (social siblings) in 2000 and she did again in 2001 (and 2002: the three-year old “Uncertain” female in Group 115 in Tables 1 and 3 [we did not know her complete dispersal history]); the nestlings of her mother and KB. In 2001, after JN had begun incubating her second clutch of eggs, FX left Group 115 and was observed intermittently working on a nest with an unmarked individual in an area at least two territories away. The nest-tree location was at the periphery of our study area (just east of Group 39; Fig. 1a) and we knew little of the details of local land ownership. After approximately two weeks, activity at this nest ceased and about one week later FX moved back in with Group 115 (and, as above, fed nestlings). FX remained with Group 115 through her disappearance in West-Nile Fall of 2002.
c. PH: We caught the male auxiliary of Group 7 (PH: the son of the breeders) in November 1998; he was at least in his second year and was likely the unmarked adult with this group the previous nesting season. He remained with Group 7 until the beginning of the nesting season of 2002, when he moved out but remained connected to group members (he was occasionally seen interacting with them and/or hanging in areas peripheral to their territory, but would also intermittently disappear for a few to many days at a time). PH’s mother was killed while incubating eggs (in a second nest) and two days later an adult female moved in; at this point the three adult auxiliaries (all previous nestlings of Pair 7; Table 1) dispersed (a female and a broodmate of unknown sex were never seen again, and their brother moved in with an older brother breeding next territory) and PH moved back in (to now Group 57). PH at first worked amicably with his father (JE) and his mate on their nest (PH made 4 of 22 nest-building trips during two watches, and was present at the nest with both breeders), but then increased attempts to get close to the female, to the point of relentlessness, in the days prior to and immediately after incubation began. He was repeatedly denied by JE. The interactions between the two males were not overtly aggressive (Appendix B Predictions 7 and 8, 2). PH stayed and cared for nestlings (he did not sire either one). The female breeder was the only one of Group 57 to survive through the fall of 2002.
d. TM: TM was marked as a nestling in Group 111 (a group no longer extant in 2001; above, 1a) in 1998; one of four broodmates in the nest of AM and XT. The following year, early in the nesting season, AM was injured during nest building and removed from the field (above, 1a) and XT nested with a replacement female (MG [Appendix C1ii]). TM remained home through the failure, a few days after hatching, of MG and XT’s first attempt (he and his three broodmates were often seen at the nest during incubation – sometimes behaving submissively but most times not, and often making variable vocalizations and behaving as if excited – and TM was seen feeding an incubating MG on several occasions) and the building of their second nest, at which point he moved next territory, into Group 10 (r with both breeders = 0 [Appendix C1iii]). Group 10 had nestlings in their nest at the time and TM fed them; we did not observe any aggression toward TM as he shifted groups. In 2000, at 2 years old, TM left Group 10 briefly and built a nest nearby with an unmarked crow, abandoned it and moved back into Group 10 (Appendix C1v), helped to raise nestlings, and remained for the following year. In 2001, during nest building, TM and a younger male member of Group 10 were observed to visit the nest of one of their neighboring groups (#115; Fig. 1a) while group members were away. TM got in and sat down briefly before both left the area. Two days later TM was seen to again visit Nest 115, this time alone and with the male breeder of Group 115 perched nearby. TM stayed on the rim for 10 sec and left (possibly because of an aborted approach by a yearling male auxiliary), at which point the male breeder briefly visited the nest. Later in the season TM again briefly left Group 10 to build a nest with an unmarked crow; incubation began (with at least one egg in the nest; observed the previous day) but the attempt was abandoned shortly thereafter, and again TM moved back into Group 10 (Appendix C1xi), helped to raise nestlings (the nest failed), and remained for the year. In 2002, Group 10 (first attempt) included four adult auxiliaries (TM and three other adults, including one that was unmarked [Appendix C1xiv]). A week or so after the female breeder of Group 10 (KT) began incubation, TM and the unmarked auxiliary began building a nest at the eastern edge of 10’s territory. The two other auxiliaries were often in the area of the latter nest over the next few days as TM and the unmarked individual continued to work. Three days after our first observation of TM working on this other nest, the breeding pair (#10) abandoned their first attempt and took over TM’s nest. At this point the unmarked individual and two other auxiliaries left the group (Appendix C1xv); the latter (of which the female was RM) appeared to attempt to nest elsewhere in Stillwater, but we lost track of them. TM remained with Pair 10, helped them complete the nest, fed KT during incubation, and fed nestlings (n=3; none fathered by TM). KT disappeared early in the fall of 2002, likely a victim of West Nile virus, at which point RM moved back into Group 10 (which included the widowed male breeder [TO], TM, and two surviving juveniles). In mid-March 2003, RM and TO began working on a nest and TO was seen thwarting TM’s attempts to carry material to it. TM left Group 10 and paired with a 3-year old female (DX, above, 1b) who, after returning home (Group 27) in 2002 and then dispersing upon the death of her mother (above, 1b), had been observed around town several times in the intervening months. DX and TM fledged three young; only one of the latter of this family of five survived the West Nile virus epidemic of fall 2003 (Caffrey et al. 2005).
e. HE: HE’s parents were unmarked and so information regarding them was unavailable, but (possibly because of land-development-related disturbance) HE and his surviving broodmate (a female) both moved south from their natal territory in late fall of their first year (1998). HE moved in with Group 19 (a group 5-6 territories away and, again, one that regularly included short- and long-term transient members; HE’s r with both breeders [female HL, male UB] = 0) by at least early March of 1999 and remained with them through early 2001, when he began spending time with a nearby group before budding a territory off from Group 19 and nesting with an initially unmarked female (despite the presence of an adult female to whom he was not related in Group 19; he had been spending time with a *different* adult female in Group 19 until she disappeared [below, 2f]). HE and his mate (Group 45; Fig. 1a) produced one fledging in 2001; a male (RZ) fathered by a male auxiliary (CO) in Group 28 (one territory away; Fig. 1a). (CO was related to UB by r = 0.6241 and was twice seen visiting with Group 19 – once near their active nest tree – during that nesting season.) RZ remained with Pair 45 through the following breeding season (2002), during which he was also seen visiting with Group 19 on several occasions (including once when foraging with UB and another group member underneath Group 19’s active nest tree). Nest 45 was not visible in 2002, and from it fledged two young that we did not catch and whose fates were unknown. HE, his mate, and RZ all disappeared by mid-October 2002.
f. DE. DE hatched in 1998. Her mother (AM; above, 1a) was injured during the nesting season of 1999 and removed from the field, and was quickly replaced by MG (Appendix C1ii). DE remained with her father (XT) and MG, helped to raise their young of the year, and moved out in November as she joined a returned AM and her new mate (Group 29, above 1a), establishing a territory adjacent to XT and MG (Appendix C1iv). In March of 2000, DE left Group 29 and was observed a few times over a few days with an adult immigrant male auxiliary (HE; above, 2e) in Group 19 (Appendix C1v), in areas along the southern periphery of 19’s territory (Fig. 1a). (Pair 19 was lazily working on their nest at the time.) Within a few more days we twice observed DE and HE arrive at an area where the breeders of 19 were foraging or loafing to have the female breeder (HL) aggressively approach and chase away DE (followed by HE). On a morning soon thereafter, DE remained back while HE approached the male breeder (UB), foraging alone on the ground. HE performed a submissive display (Verbeek and Caffrey 2002), something we had possibly not seen between these two since early in the period after which HE originally moved in with Group 19 (in early 1999; HE was not yet one-year old at the time). In field notes (CC): “Could he be asking permission for DE to stay?” Over the next three weeks, as Pair 19 completed their nest and incubation, DE was seen in closer and closer proximity to UB and then HL (when off the nest), with few to eventually no more threats against her. DE was never seen at Nest 19 but was often in its general area with other group members, especially HE. The special relationship she had with HE – they spent a lot of time together alone – lasted through the end of December (2000), after which DE was never seen again. As above (2e), in the Spring of 2001, HE budded off a territory from that of Group 19 and bred with a female of unknown origin.
g. GN: Groups 6 and 19 (Fig. 1) were next-territory neighbors whose members were regularly seen together. During the breeding season of 2000, Pair 6 was assisted in nesting by three yearling males (broodmates: GN, DQ, and OV). DQ moved in with Group 19 (r with both breeders [female HL, male UB] essentially zero) in the fall of 2000; Group 19 already included two adult male auxiliaries (HE [an immigrant; above, 2e] and OZ [of uncertain relationship to HL and UB but whose r values with them indicate they were related]), two adult female auxiliaries (a daughter of HL and UB, and an immigrant [DE; above, 1a and 2f]), and one hatch-year female (daughter of HL and UB). As the breeding season of 2001 got underway, OV budded a territory off from Group 6 and bred with an unmarked female (Group 50, Fig, 1a; OV was the only member of our population to attempt to breed independently in his first year of maturity, at three years old), and HE, OZ, DQ, and DE moved out of Group 19. GN had twice been observed spending time with DQ (with Group 19) in the weeks before DQ moved out of 19, and a few weeks after the departure of HE, OZ, DQ, and DE, and after incubation had begun at Nest 19, GN joined Group 19 in addition to maintaining his membership in Group 6; he remained a member of both groups through the end of the summer of 2001, after which he moved out of Group 6. Early in the breeding season of 2002, GN was observed working on a nest with HL; at a nest-building watch done the following day, GN made 8 of 13 nest-building trips and UB did not go to the nest (Caffrey et al. 2015a). (UB had a leg injury that intermittently worsened and improved over the years, and he was noticeably pained and weakened during the nesting season of 2002 [and 2001].) Of 40 total nest-building trips during four watches over seven days, GN made 22, HL made 15, and UB three (Caffrey et al. 2015a). Work on the nest then stopped, and members of Group 19 were not seen at the nest or in the nest tree again until 10 days later. During this period HL, UB, and GN (and two other group members) were observed several times in other areas of their territory, but on many occasions could not be found (we thought this nest had been abandoned). Over the subsequent six days (prior to the start of incubation), the pair and GN made infrequent, brief trips to the nest during our nest checks. During incubation GN fed HL and was present at the nest more often and for more time than UB (Caffrey et al. 2015b). GN made four of four nestling feeding trips during watches on the first and second days of hatching. We caught him the next day to replace a missing patagial tag, at which point he moved out of Group 19 and partially out of our population, and partially back in with Group 6 (where he made at least one nestling feeding trip; difficult viewing conditions prohibited detailed observations). He was observed to visit Group 19 twice before partially moving back in one month later (on one visit he made at least one nestling feeding trip), and thereafter he regularly spent time with both groups (19 and 6) through October, when he and several members of Groups 19 and 6 disappeared, likely the victims of West Nile virus. The single nestling in Nest 19 in 2002, which fledged successfully, was the daughter of HL and UB.
h. KB: KB was caught with Group 14 (r with both breeders = 0) in early December 1998; he was likely in his second year. He remained with Group 14 through the end of January 1999, at which point he moved in with Group 18 (a group west of Group 14 [Fig. 1]; r with breeders = 0.1122 [female] and 0.183 [male]). In early April (1999) he was again seen with Group 14; through the end of that year he was a member of both groups 14 and 18. Early in 2000 KB ended his association with Group 14 and was seen with, in addition to Group 18, Group 15 (r with female breeder [JN] = 0.1148, with male [ED] = 0); he then moved in with the latter and ultimately assisted in feeding JN and ED’s nestlings. Over the course of the breeding season and summer of 2000, ED suffered increasingly from an injury to one leg, and he disappeared by the end of the year. KB remained and bred successfully with JN (Group 115) in 2001 and 2002 (and 2003; both survived the fall of 2002).
i. TF: TF was caught in November 1998 with her mother (HL) and her mother’s mate (SF; Group 12), and another second-year female. SF was not TF’s father (her father was alive and the current breeder in Group 27) but *was* the father of the other second-year female (BN; not HL’s daughter). TF and BN remained in Group 12 through 1999 and 2000. In May 2000, a yearling male (AK) moved in with them. BN moved out of Group 12 in January of 2001, and in early February, TF and AK were seen several times together in the area bordering the territories of Groups 12 and 14 (Fig. 1a) before AK moved out (mid-February). About two weeks later an unmarked adult (“Um”) moved in. TF and Um were observed several times alone together in early-mid March in the same area where she and AK had spent time. On one occasion in mid-March, TF was foraging together with Pair 12 (while one of their yearlings repeatedly harassed – ran at -a flock of resting gulls in the background) when the female – TF’s mother – was observed to approach TF a few times in ways that made TF hop back a step or two before resuming foraging. The approaches by the mother appeared lackadaisical and were followed immediately by everyone resuming normal behavior. TF was seen with Um a few more times before she moved out at the end of March (she was seen several times in areas surrounding Group 12’s territory but never near the nest or interacting with the pair, until September, when she moved back in). Um, on the other hand, remained with Group 12, was often with them, and was observed once with a full esophageal pouch, once landing uncontested on the nest rim, and once seemingly attempting to sit in the nest with HL. In April, AK was seen visiting with Group 12, and in July a female in her third year (BM) partially moved in for approximately two months (she seemed to also spend time with neighboring Group 44 [Fig. 1; not included in our study because of difficult viewing conditions]; she was an immigrant in both [she was the daughter of Pair 11; above, 1a]). In early 2002, TF and Um were again seen alone together several times at the southern end of Group 12’s territory, once in a calling/swooping battle with Group 14 (Fig. 1b). BM joined Group 12 again in February and was seen by herself a few times, in the area visited by TF and Um, repeatedly producing vocalizations we associate with territory owner-ship – we wondered if she was attempting to set up a territory on her own – as TF and Um (caught soon thereafter and tagged EC; Group 40) instead budded west across the lake and set up a territory in some available habitat plus some of what had been Group 4’s southeastern end (Fig. 1a and b; Group 4 shifted some of their activity west as groups 40 and 55 formed). Both TF and EC survived through the fall of 2002; their two young of the year were not so lucky.
j. EK: Early in the summer of 2000, EK (above, 1a) was one of three surviving broodmates from Nest 11, the nest of MG and XT (above, 1a and Appendix C1vi). Group 11 at the time also included two of EK’s two-year old half-brothers (NK and CS; a third, TM was living with the next-territory group [above, 1a and 2d]) and a two-year old half-sister (DE; above, 1a and 2f). MG disappeared in June (Appendix C1vi) and YL, an adult female, moved in in October (above, 1a and Appendix C1viii). In early 2001, NK ended up pairing with YL (Group 42) and taking over ownership of his natal territory from his father (above, 1a and Appendix C1x). EK and his broodmate brother (ZO) remained and helped to raise YL and NK’s offspring (his broodmate sister [AI] moved out at hatching [above, 1a, below, 3f3, Discussion, and Appendix C1xi]). Although EK was presumably not capable of cuckoldry (he was a yearling), NK was observed six times throughout the nesting period to act dominantly toward EK, by displacing him from the nest area or from food (NK was never observed to act this way toward ZO). Most of the time, however, nothing appeared amiss between NK and EK; EK made many trips to the nest over the season carrying nesting material or food for YL or nestlings. Especially during the first part of the nestling stage, EK often spent long periods of time (maximum observed bout = 14 minutes) gazing down into the nest and (we presume) gently preening nestlings. NK was twice observed passing food to EK to feed nestlings, and twice to allow EK to approach (on the ground) and take some of the food he was eating (the contents of an avian egg, and a peanut; on the latter occasion, EK begged upon approach and was fed by NK). ZO left Group 42 around the end of 2001 (Appendix C1xiii) but EK remained and, early in 2002 -before YL and NK began working on their nest – began a nest with a female auxiliary from Group 7 (RA, Fig. 1b). Both EK and RA maintained pre-hatch membership in their respective groups, through abandonment of their nesting effort, and resumed complete (pre-hatch) residence with those groups thereafter (Appendix C1xiv). EK amicably associated with YL and NK (and their one-year old son, SN) through nest building in 2002, contributing a few times (during one nest-building trip, after working in the stick he had brought, he sat in the nest and adjusted material for nine minutes). All four members of Group 42 disappeared for a few days after work on the nest ceased. Upon their return, EK was being actively denied access to YL by NK; he could not land anywhere near YL when she was away from the nest or anywhere near the nest without being chased by NK (but when YL was not involved, NK and EK behaved normally toward each other). This continued through incubation. Once the young in Nest 42 hatched, EK was welcome again and was a regular feeder of nestlings. NK disappeared by mid-September, one of the first of our population to die of West-Nile-virus infection (Caffrey et al. 2005), and in the three weeks before EK then disappeared, too, he was with YL in 100% of our observations. YL and NK’s two young of the year disappeared that fall as well, and YL bred with SN – her 2-year old son – in 2003 (CC, unpubl. data).

#### 3. Pre-hatch auxiliary dispersal behavior

Of 18 Pre-hatch auxiliaries that dispersed, one was unmarked and so could not be followed, and three were never seen again subsequent to their disappearances from groups.

a. Four individuals (KP and PR [Group 333, 2001; above, 1b], RA [Group 7, 2001], and LH [Group 27, 2002; above, 1b]) were seen regularly at the peripheries of their groups and in surrounding areas – floaters considered to be “highly philopatric dispersers” (Penteriani et al. 2011) -until moving back in later in the year (RA was known to visit occasionally during the intervening months).
b. Two individuals visited during the nestling stage before they were never seen again:

1. A one-year old female (BO) returned for a brief visit three weeks after her dispersal (she fed nestlings twice during one watch).
2. A 3-year old immigrant male (RG; above, 1b) returned to visit several times during the nestling stage. During six of 19 nest watches, his first visit being after two weeks post-hatching, he was seen to make three feeding trips and five nest visits, the latter in which he simply looked down at nestlings (all while the female was temporarily away).
c. One 2-year old female (LH: Group 333, 2001; above, 1b) began her move across town into a group (#27) in which she ultimately became the breeder.
d. One adult male and female (RM) from the same group (#10 in 2002) dispersed and thereafter appeared to work on a nest within our study population area (we lost track of them). RM returned to Group 10 later in the year (to ultimately pair with the widowed male breeder (above, 2d).
e. One adult male (KP) left his group (333 in 2002) at hatching and nested with an unmarked crow about 150m away from Nest 333; he moved back in with Group 333 three and a half weeks later, after he and his incubating partner abandoned their attempt (above, 1b).
f. Three individuals were known to have moved out of our study area:

1. An adult male was seen occasionally in foraging flocks outside of Stillwater (this individual moved back home late in the year).
2. An adult female (SV) who dispersed in 2001 (above, 1b) occasionally returned to Stillwater and visited with her family. She was seen to feed nestlings on one such visit, which lasted about four days, in 2002.
3. The 1-year old female (AI) that moved to Pawnee, OK (above, 1a, and Discussion). Interestingly, in retrospect, in the field notes from the nest watch when AI was last seen is reference to the unusual “Lots of calling this a.m. as if keeping in touch with one another.”)
g. An adult immigrant male (GN) left his group (#19) likely in response our disturbance (above, 2g). He moved partially out of our population and partially back in to his natal territory (Group 6) where he fed nestlings; after a month and including at least two visits (including a nestling-feeding trip), he moved partially back in with Group 19 and thereafter maintained partial residence with both Groups 19 and 6.

#### 4. Extended familial care

Regrettably, many nestlings in 1999 were marked with patagial tags that ended up causing high mortality and harm to those that survived fledging and whose tags did not fall out on their own (Caffrey 2002b). Many hours were spent in attempts to rescue burdened individuals and although several were saved, a few resisted capture and continued to decline in body condition until they disappeared. One such individual, a female (TN, in Group 19 [above, 2g]), could no longer fly by the beginning of February 2000 and was restricted to a small, dense patch of woods in a residential area. Her father (UB) and sister (a broodmate) would return to the woods from other parts of their territory regularly (at least a couple of times a day) and spend time near her. One of us (CC) observed UB feeding TN twice in late February, before she disappeared.

## APPENDIX B.

### Emlen’s Predictions

In a seminal paper, Emlen (1995) proffered 15 predictions regarding group formation, group dynamics associated with breeder deaths and replacements, within-group aggression, and the sharing of reproduction within groups. Here, in the contexts of those predictions, in acknowledgment of the many axes along which individual animals may differ in behavioral phenotypic traits, and supported by recent demonstration of *Corvus* awareness of their surroundings and the relationships among the contents thereof (e.g., Taylor et al. 2007) -including the intentions, and emotions, of conspecifics (e.g., Bugnyar 2013, Bugnyar and Kotrschal 2002, 2004, Bugnyar and Heinrich 2006, and Fraser and Bugnyar 2010b) - and their ability to plan for the future (e.g., Dufour et al. 2012), we risk anthropocentrism to imagine some of the points of view of individual crows in Stillwater. We saw only small parts of their lives, and were almost completely unaware of how individuals spent their time, and with whom they interacted, while not within our 6.4 x 9.5 km study area (Fig. 1), including where, and with whom, they spent their nights. Yet we knew our marked individuals to have different habits and different personalities (Caffrey et al. 2015a), and we knew they had social relationships of varying quality (Fraser and Bugnyar 2010a) with other crows in their own and other groups, and with those floating in and out of the population.

**Prediction 2** of Emlen (1995) has to do with the fact that high quality resources are generally worth more than those of lower quality, and for individuals choosing between leaving natal situations or staying, the options available once leaving would have to be of high worth to justify dispersal out of high-quality natal situations. As such, prolonged philopatry on high-quality territories leads, through inheritance, to “dynasties” (Emlen 1995): generations of single lineages continuously occupying the same areas. Our study did not span more than a single crow generation (in years), yet we observed responses to deaths of breed-ers -the necessary first steps to inheritance.

Unlike Emlen predicted, and more in line with the norm of more recent studies (Ekman et al. 2004, Dickinson and Hatchwell 2004), territorial inheritance in our population was rare. Except during the West-Nile-virus years of 2002 and 2003, only single members of pairs died/disappeared at a time, and widowed females were not evicted (below, Predictions 7 and 8). Surviving breeders of both sexes paired with new mates, and thus much of the space in Stillwater was continuously occupied by pairs consisting of survivors and replacements, the latter then sometimes becoming the survivors. We did not observe aggressive behavior by or toward anyone in the context of breeder replacement (below, Predictions 7 and 8), and in only one instance of nine breeder deaths/ disappearances during the years of our study (five males and four females, including AM’s removal from the field; Appendix A 1a) did the surviving breeder not maintain ownership of his territory. In that case, the widower (XT) left as his son (NK) paired with a replacement for his step-mother (Appendix A 1a), and ended up pairing with his previous mate (AM) – NK’s mother – in an adjacent territory (after the death, shortly after XT left his territory, of AM’s [then] current mate [Appendix A 1a]).

**→** Might XT have been “hoping” for an opportunity with AM and so didn’t much contest his son’s “usurption?” (Because he didn’t: Appendix 1a.)
**→** What role might the behavioral compatibility of mates (Ihle et al. 2015) play in such decisions?

**Prediction 5** contends that under usual conditions, sexual aggression within groups of relatives should be lower than in “otherwise comparable groups composed of nonrelatives” (although the latter are not discussed; Emlen 1995), because of incest avoidance; sons will rarely compete with fathers, or daughters with their mothers, for sexual access to their parent of opposite sex (Emlen 1995). In keeping with the prediction, except for in 2003 (in the aftermath of the West-Nile-virus epidemic of late summer and fall 2002), there was no evidence of incestuous matings among our population members, despite ample opportunities to pursue such unions, as is the case for most cooperatively-breeding birds (Koenig and Haydock 2004). However, in contrast to the prediction, there was not heightened sexual aggression in groups with members unrelated to breeders. We did not see much evidence of aggression in our population at all (which is not to say that “hidden threats” of eviction, departure, and attack (Cant 2011) were not maintaining the peace): in 2001, for example, in detailed field notes documenting thousands of person-hours of observation, in addition to the six times NK advanced toward or displaced EK (Appendix A 2j), we recorded a total of only seven occurrences of dominance behavior toward other individuals within focal study groups (six occurrences involving individuals of unknown identity [e.g., in groups with more than one unmarked member] were also recorded). All of the seven incidents occurred while participants were foraging, all but one occurred during the months February-April, three prompted submissive displays by targets and four their simply moving back a bit (including TF; Appendix A 2i), only one involved an immigrant, and all seven were followed by immediate resumption of previ-ous activities.

**→** We wonder why immigrants, particularly males, did *not* attempt to reproduce within their groups, and the few explanations posited by others (references in Cockburn 2004), e.g., those having to do with not jeopardizing relationships with male breeders because of their “main provisioner” status, do not seem relevant.
**→** Could “holding back” perhaps have been a form of “paying to stay” (Gaston 1978), given all of the benefits to group membership (Discussion)?
**→** How might individuals have decided/ settled such things?

**Predictions 7 and 8** both have to do with within-family conflict in the context of breeder death and mate replacement.

**Prediction 7**: that group members will compete among themselves for resource ownership/ breeding status and that, because of incest avoidance and the fact that males usually dominate over females (Emlen 1995), widowed females will often be evicted by older sons (whereas the same is not the case for widowed males and their daughters).

**Prediction 8**: that once a parent has re-paired, same-sex offspring should compete with them for access to the replacement, and the intensity of the aggression between competitors (fathers and sons) should increase with decreasing asymmetry in their dominance status (Emlen 1995).

Five of five female breeders widowed between Jan 2000 and May 2002 were not evicted from territories. One female had no potential competitors present and in one case we did not have complete pre-replacement information, but three cases were in groups with adult male auxiliaries present: a son in each and also in one an immigrant (the immigrant paired and ultimately bred with the widowed female [KB; Appendix A 2h]).

In addition, we reiterate that we saw little aggression within groups, including in the following situations, predicted to elicit it:

1. We had been observing Group 3 for more than a year before we had one of the two adult males marked, and it wasn’t until four months later (March 1999), when the marked one (SW) was the one helping the female build the nest, that we thought we could distinguish breeder from auxiliary (NX); prior to that both seemed to behave as breeders and the two were completely at ease with each other. As it turned out, NX fathered two of Group 3’s four fledglings that year (Appendix A 1b), likely a case of shared reproduction (below, Predictions 12-15). In March 2000, the female breeder disappeared and a replacement moved in. (This was preceded in January by the immigration into Group 3 by an adult female [PR; Appendix A 1b], with whom NX ultimately paired [in 2002] and moved into a budded territory.) It was nesting time when the replacement (DZ) moved in and, as described in Appendix 1b, it appeared to us that SW was pairing with DZ (creating Group 333), yet NX was often with them or close by while they foraged or loafed. NX fathered one of the two fledglings (of three) for which we had DNA samples that year. NX remained in Group 333 through another year and a half before moving out with PR, and infrequently fed nestlings and fledglings in 2001; he was not the father of any of the three fledglings but was the grandfather of one (Appendix A 1b). Only once between February 1998 and May 2002 did we ever see any evidence of status establishment by SW or NX; an occasion involving both strutting with ruffled feathers before resumption of foraging.
2. PH was an adult male auxiliary, son of both breeders, and at least five years old when he moved out of his territory in March 2002. He continued to occasionally interact with his former group (#7) at territory peripheries but would also disappear for days at a time (Appendix A 2c). Upon his mother’s death – during incubation late in the season – and the moving in of a replacement, PH moved back in and participated in nest building, although his contributions were constrained by his mate-guarding father (JE): with increasing intensity as nest completion and incubation approached, PH continually attempted to get close to JE’s mate and JE continually denied him access. Their interactions never involved overt aggression; JE simply repeatedly put himself between PH and the female, and PH simply repeatedly left the immediate area temporarily, to return to her whereabouts shortly thereafter. Situations 3 and 4 do not involve breeder replacements but are cases in which the reproductive interests of brothers were pitted against each other:
3. RN was turning four years old when he moved to an adjacent territory (in January 2000) to pair with a (presumably widowed) female (creating Group 33); his turning-two-year-old brother (KR) moved with him (Appendix A 2a). KR remained with RN and his mate (GP) through the next two and a half years, participated in nesting activities in 2001 and 2002, and was never seen attempting to reproduce with GP; nor was RN ever observed to aggress against KR (in fact, Group 33 was an especially close-knit group; Appendix A 2a).
4. NK was the son who acquired ownership of his natal territory from his father early in the nesting season of 2001 (above, Prediction 2, and Appendix A 1a). EK, one of NK’s yearling brothers, opted to stay and help raise nestlings, as did the other (ZO); their yearling sister moved out at hatching (Discussion, and Appendix A 3f3). EK chose to remain with NK and his mate (YL; Group 42) through the next year, as well (ZO did not). As was the case with PH (above, number 2, and Appendix A 2c), although EK was actively kept away from YL during her fertile period in 2002 (Appendix A 2j), the interactions between NK and EK involved only displacements and were never overtly aggressive (although a year earlier we had observed six instances of dominance behavior by NK toward EK; Appendix A 2j). **→** Might the difference in KR’s and EK’s behavior have had anything to do with the differences in their relationships to their brothers? (KR was a full sibling of RN; EK a half-sib of NK’s.) If so, why?
**→** Regarding the instances of dominance behavior of NK toward EK but not toward ZO (another yearling half-brother group member), did NK “know” that EK would remain through sexual maturity, and would thus eventually pose a reproductive threat, but ZO would not? How?

**Prediction 9** contends that replacement breeders should not only *not* invest in existing offspring but might also consider killing them (Emlen 1995). If investment includes the sharing of resources, then mate replacements *did* invest in existing, independent offspring. With regard to dependent offspring unrelated to replacements, our two instances are confounded by genetic and social relationships:

1. XT was the father who left his territory to his son and mate and then – weeks later – moved into the adjacent territory with his previous mate (AM), after the death of her current mate (MZ) and during the late nestling stage at her nest (above, Prediction 2, and Appendix A 1a). XT was observed to feed the single remaining nestling during two final nest watches (Appendix A 1a). The nestling was the offspring of AM and MZ, and XT was related to MZ by r=0.5207 (Appendix A 1a), and so he was feeding a close relative. But we were providing AM with food at that point (Appendix A 1a), and so presumably XT’s “help” was not necessary for nestling survivorship.
2. It was six days post-hatching in Nest 333 in 2002 when the male breeder (SW) disappeared. KP, an adult male pre-hatch auxiliary (not the son of either breeder; Appendix A 1b) who had sired at least one of the nestlings in Nest 333 (but none of the surviving fledglings), had moved out of the group at hatching. He returned to Group 333 upon abandonment at his own nest (Appendix A 1b; 2.5 weeks after the disappearance of SW), was not aggressive toward anyone, and although not subsequently observed to feed nestlings or fledglings, was suspected to have done so based on circumstantial evidence (possessing a full esophageal pouch, and sitting with begging fledglings).

**→** As there was no bringing their new mates into new reproductive cycles through killing their dependent offspring (and again, nothwithstanding the relat-edness between XT and the nestling he fed), it appears these males may have been acting to win over the interests of territory-owning females (who they knew through prior relationships, and with whom they bred the following year), possibly also contributing to increasing the lifespans of preferred partners by reducing their current parental care costs.

**Prediction 11**, which derives from considering only the indirect benefits available to auxiliaries through helping and the direct benefits of mating with replacements, has it that families with replacements will be less stable: that auxiliaries will be more likely to disperse upon replacement of a parent (Emlen 1995). Likely because of the many possible direct benefits to auxiliary crows in Stillwater, no matter their genetic relatedness to breeders (above, Discussion), groups did not dissolve upon breeder replacement. Nor did auxiliaries of the same sex as replacements leave more often than not: for groups with complete information available, at the time of replacement of four male breeders, all auxiliaries remained in their groups (three adult males, three adult females, and three immature males). At the time of replacement of four females, both of two adult female auxiliaries left, but one was an immigrant who likely would have left anyway (Results). One of four adult males left, and all immature auxiliaries remained (three females and seven males). Except for the unusual case of Group 3/333 (below, Predictions 12-15, and Appendix A 1b), auxiliaries did not mate within groups, and as such, Prediction 11 did not hold for our population.

**Predictions 12-15** have to do with the sharing of reproduction among group members, giving rise to extended families (Emlen 1995). We revisit Group 3/333 (Appendix A 1b) briefly so as to assess the applicability of the four parameters suggested to set the conditions for, and influence the magnitude of, reproductive skew (Emlen 1995).

As above (Predictions 7 and 8, #1), we could not at first distinguish between “breeder” (SW) and “auxiliary” (NX) by way of their behavior. They were related to each other (Appendix A 1b) but were not father and son, and were likely not brothers. They lived in a group with a single breeding female, replaced once during our study (and so Group 3 changed to 333; above, Predictions 7 and 8, and Appendix A 1b), and variable numbers of other auxiliaries (up to 10 at a time), including at least one adult immigrant female (PR; Appendix A 1b), in a large (Fig. 1), heterogeneous territory. NX was a member of Group 3/333 from at least the fall of 1997 through February 2002, and contributed to nesting attempts, albeit minimally, in all years. He sired two of the group’s four fledglings in 1999 and one of the two of known parentage (three total) in 2000. KP, one of NX’s sons from 1999, stayed through fall 2000 and then moved to an adjacent territory for 3-4 months before returning to 333 for about six weeks (in early 2001) before he moved out again as hatching at Nest 333 was imminent (Appendix A 1b). Of the three young fledged from Nest 333 that year, one was KP’s and none were NX’s, and yet KP left for the nestling stage while NX remained (and contributed to feeding).

Again, we saw no evidence of aggression among these group members.

**→** What might SW have possibly gotten out of sharing reproduction, especially with a distant relative? SW disappeared in 2002, six days after hatching began at his nest; might he have sensed his impending demise and wanted a relative to inherit his productive territory? Why pay the extra costs of nest building, and any associated with being the territory owner (references in Lenda et al. 2012), if he might have been just as successful in the role of an “auxiliary?”
**→** Might NX have already known in the Spring of 2001 – after PR had temporarily moved out – that he intended to bud off a territory the following year (with PR, subsequent to her return) and was negotiating space with SW (and the female breeder of 333) through feeding nestlings (none of which were his own though one was his granddaughter)?
**→** Once SW knew NX was moving out, did he then share reproduction with KP (a more distant relative, but still a relative) so that maybe he’d stay close, so as to inherit SW’s territory? If so, it worked.

Emlen’s parameters – the genetic and social aspects of group dynamics influencing the distribution of direct reproductive benefits (among same-sex individuals) – include the potential benefits to dominant individuals if subordinates decide to stay, the potential benefits to subordinates choosing to leave, the degree of asymmetry to competitors’ dominance relationships, and the genetic relatedness of participants (Emlen 1995). Implicit in Predictions 12-15 is that dominants will only share if it is to their advantage somehow, i.e., they must be benefitting by way of subordinate presence. As such, Prediction 12 contends that as the quality of a subordinate’s options elsewhere increase, dominants should be willing to share reproduction. We had no evidence that breeding space was limiting for crows in Stillwater (Discussion) -in fact NX himself ended up budding – and there were many unpaired adult females in our population. It would thus appear that NX was biding his time, in a reproductively successful way, until he met someone with whom he wanted to pair (there are fitness benefits to be gained by choosing a compatible mate [Ihle et al 2015], perhaps especially in cases of long-term pair bonds [Gabriel and Black 2011]) and the time was right.

Given NX’s decision to stay -given SW’s ostensible sharing -Prediction 13 would have SW and NX sharing relatively equally because of the lack of difference in their dominance status, which is what we found in the two years for which we had data. There was a greater asymmetry to the relationship between SW and KP, but because if sharing is to occur the sub-ordinate has to father at least one of the young in the nest, the difference in the number of offspring fathered by the two individuals in a single nest is constrained (KP sired one of three late-stage nestlings; all three fledged). KP’s daughter was of similar size to her brood-mates but did not survive the fledgling period, and KP returned (after having left temporarily) to pair with the mother of his daughter, in his natal territory, no longer owned by either of his parents.

Our single instance of reproduction being shared approximately equally between relatives of uncertain relationships does not lend itself to addressing Prediction 14, which contrasts sharing between siblings and between parents and offspring, contending that the latter should be less equitable because of the greater difference in relatedness to offspring of the other in parent-offspring sharings. Yet as all dominants choosing to share are relinquishing direct benefits, and the indirect benefits for which they are trading are of equal value (grandoffspring versus nieces and nephews: r with both, on average = 0.25), the logic supporting Prediction 14 is not straightforward.

Prediction 15 is predicated on the notion that auxiliaries automatically “help,” and thus automatically benefit indirectly. As such, auxiliaries more closely related to breeders should require less incentive to stay, and so reproduction should be shared most with those least closely related. Not all auxiliaries in our population contributed to nesting attempts, and we have no evidence that group size was positively related to number of offspring fledged per attempt (Caffrey et al. 2015a). That, in addition to having only a single occurrence of shared reproduction, renders this prediction unassessable with our data.

### Conclusions

The predictions of Emlen (1995) having to do with within-group aggression, group dynamics associated with breeder deaths and replacements, and the sharing of reproduction were not well supported in our study population. Neither are they well supported in humans (Davis and Daly 1997), for reasons with a lot in common with crows: the social bonds shared among members of groups were strong, and groups did not dissolve upon the opening up of opportunities for nonbreeders. Crows regularly visited with former groups, and with former group members and friends in other groups. In the future, for theories regarding group formation and maintenance in cooperatively-breeding vertebrates to be applicable to crows (and humans), they will need to incorporate some of the complex psy-chological adaptations that influence relation-ships among population members (Davis and Daly 1997).

## Appendix C.

### Schematics of Appendix A 1a and b

The diagrams included here were part of a Powerpoint presentation given at the 2015 joint meeting of the American Ornithologist’s Union and the Cooper Ornithological Society by CC.

In all diagrams, as in 1ii, individual crows are indicated by their two-letter “names” (Methods).

Within groups, as in 1ii: females are on the left, males on the right (individuals of unknown sex in the center), and lines separate breeders and auxiliaries of different reproductive classes.

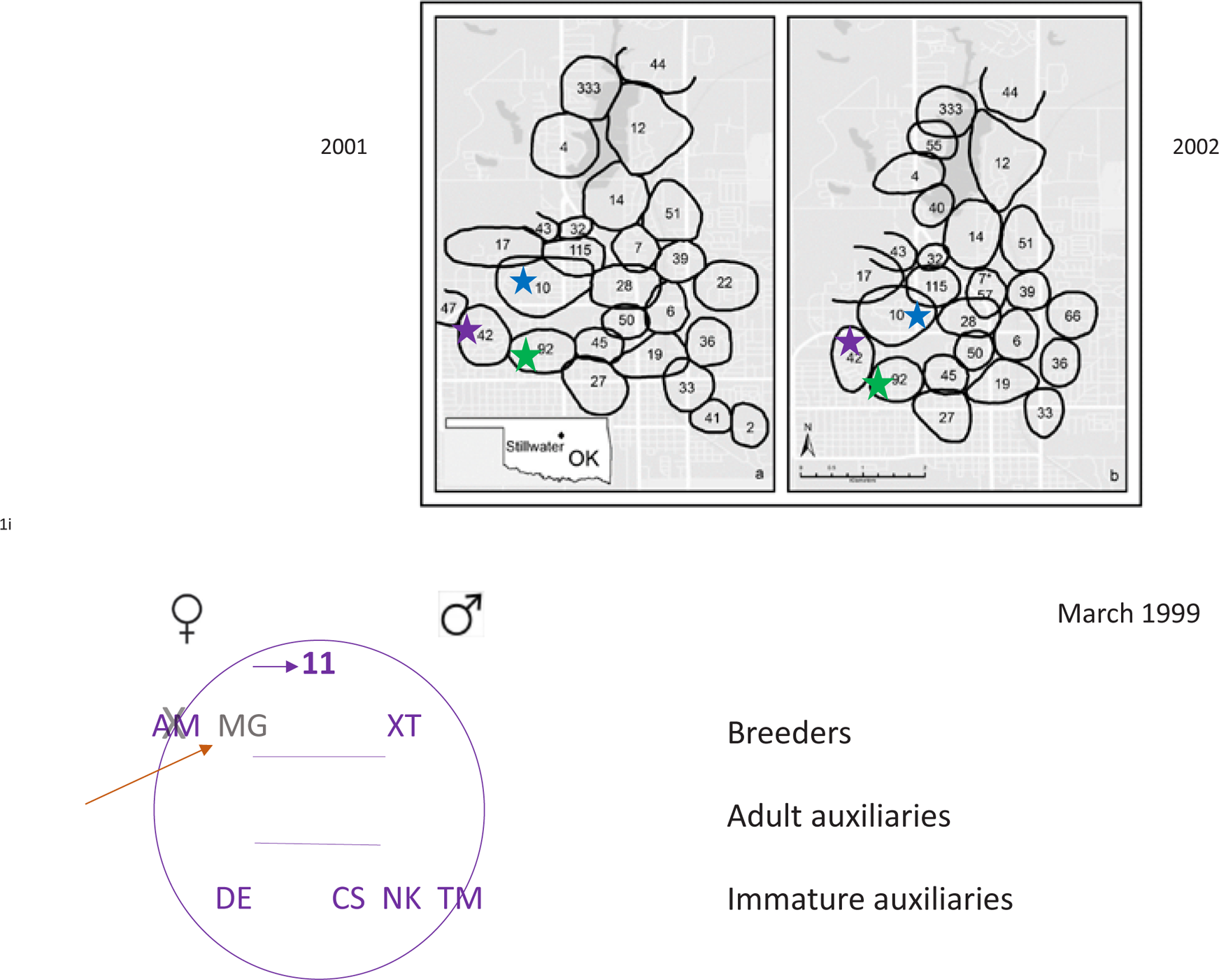
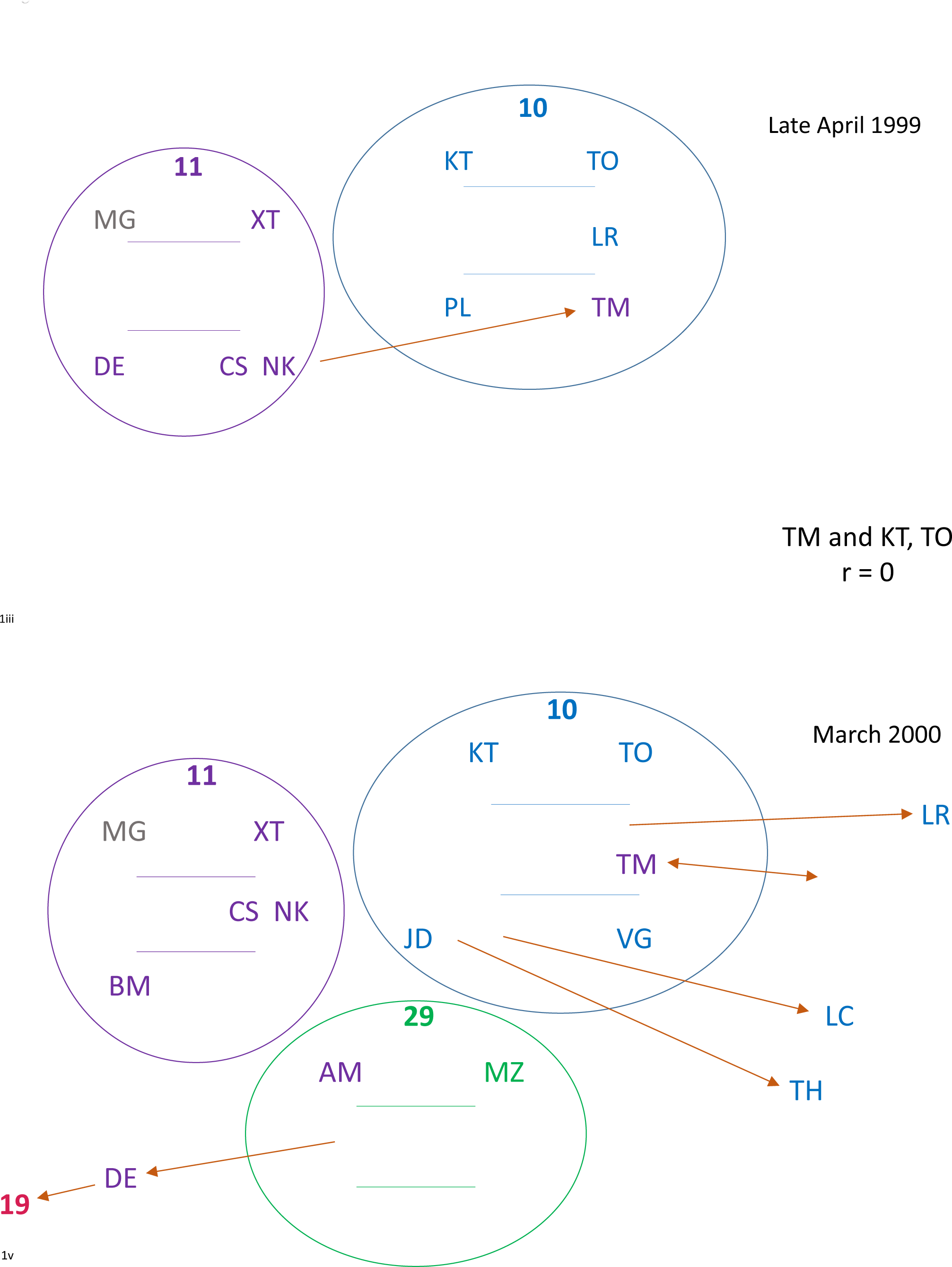
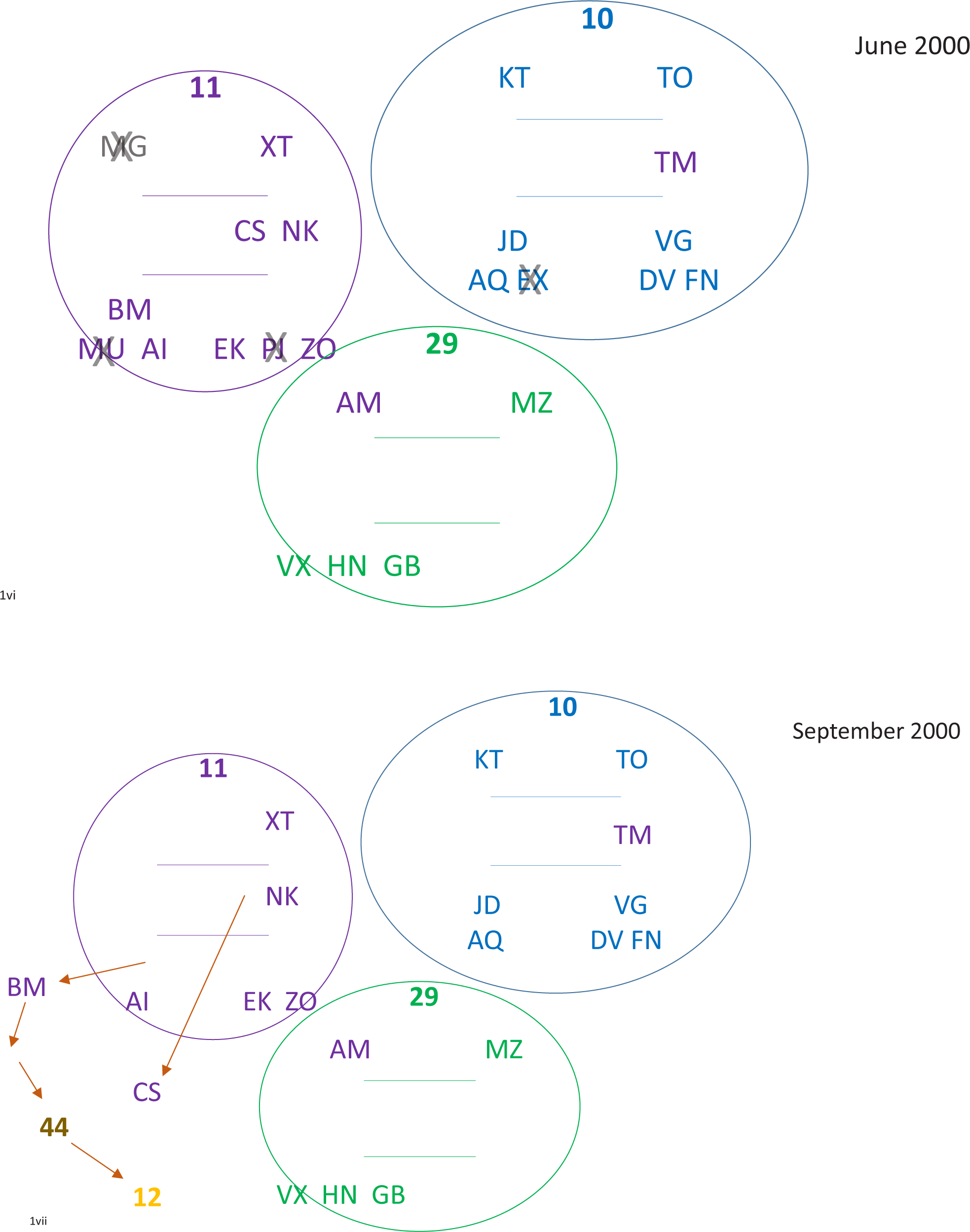
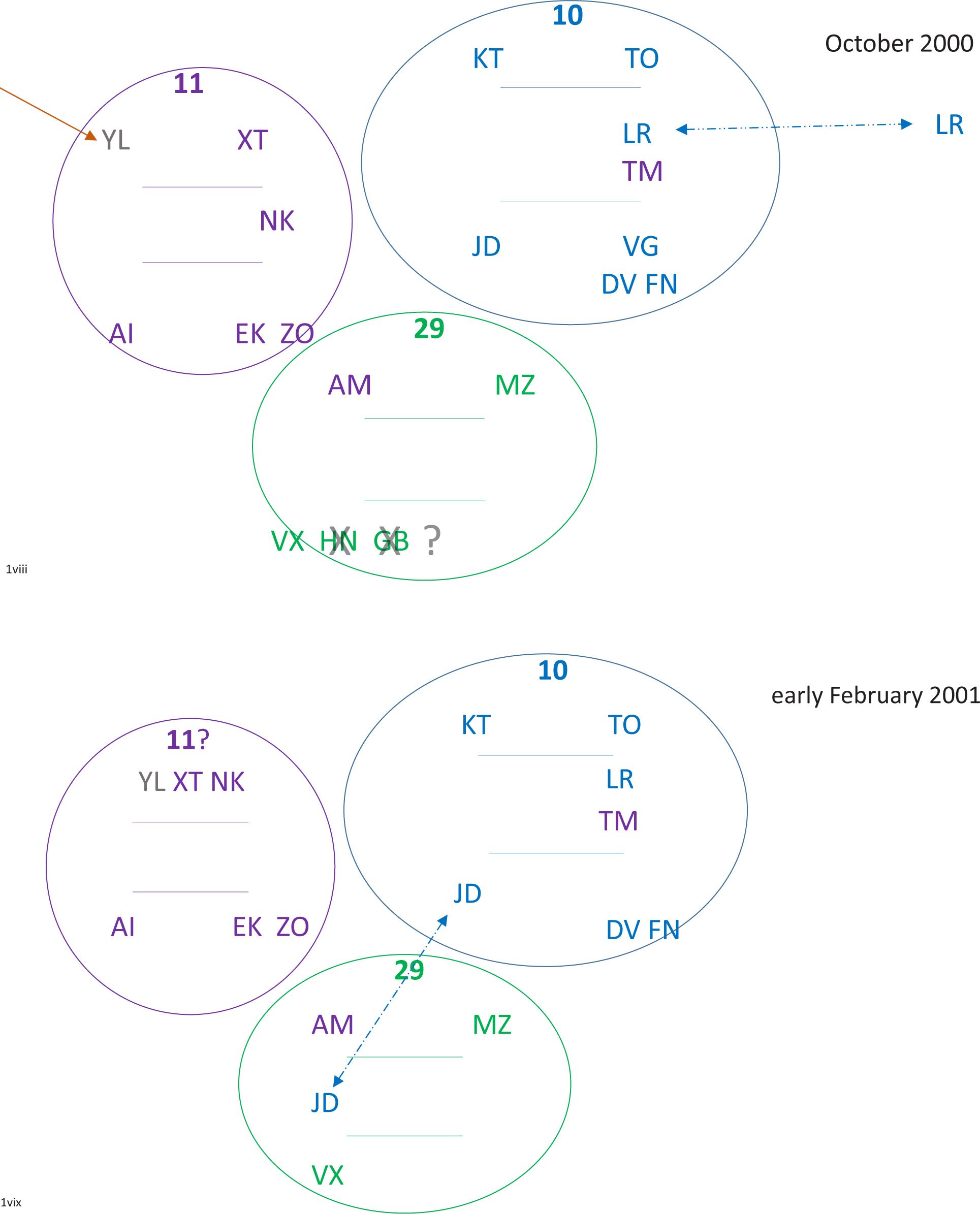
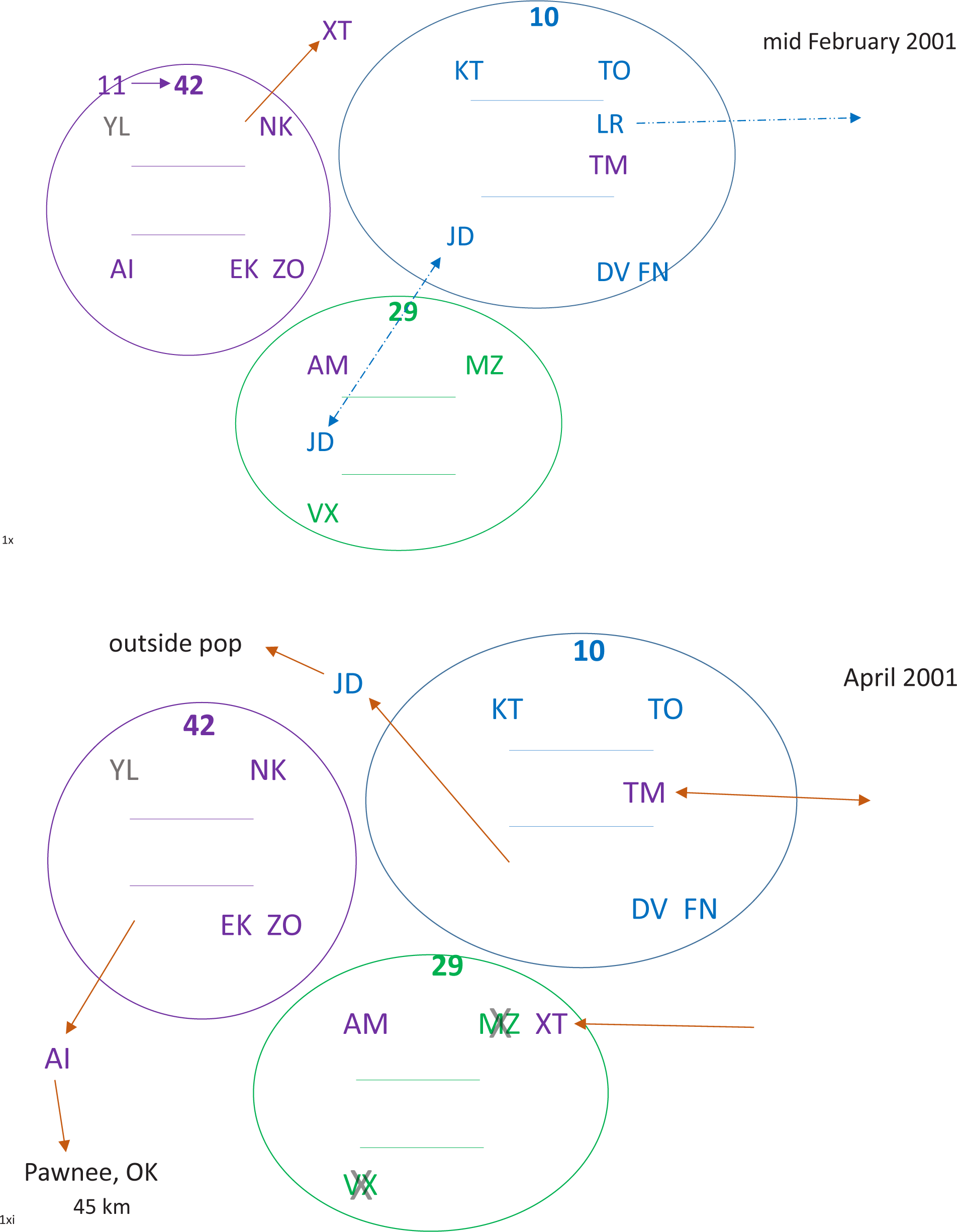
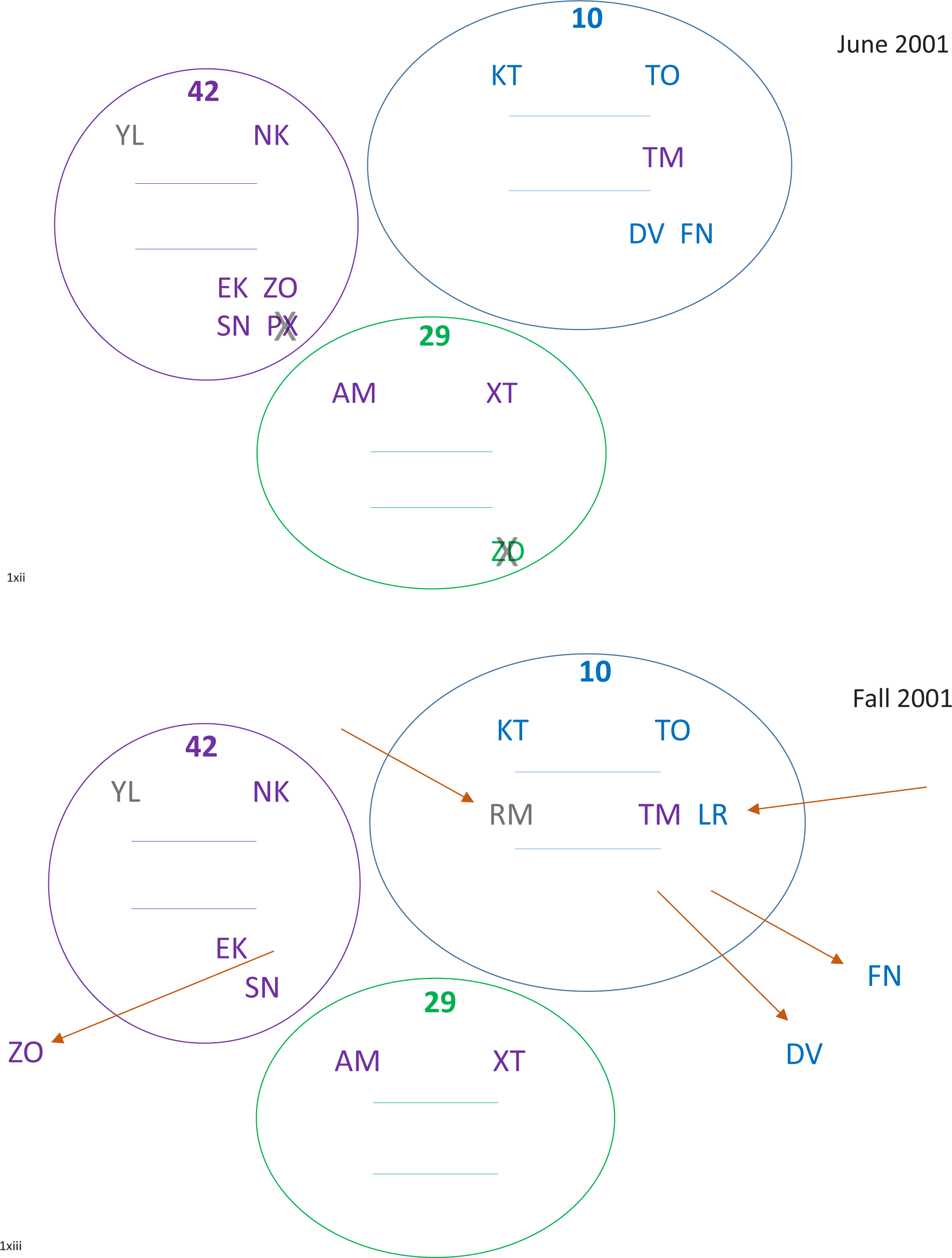
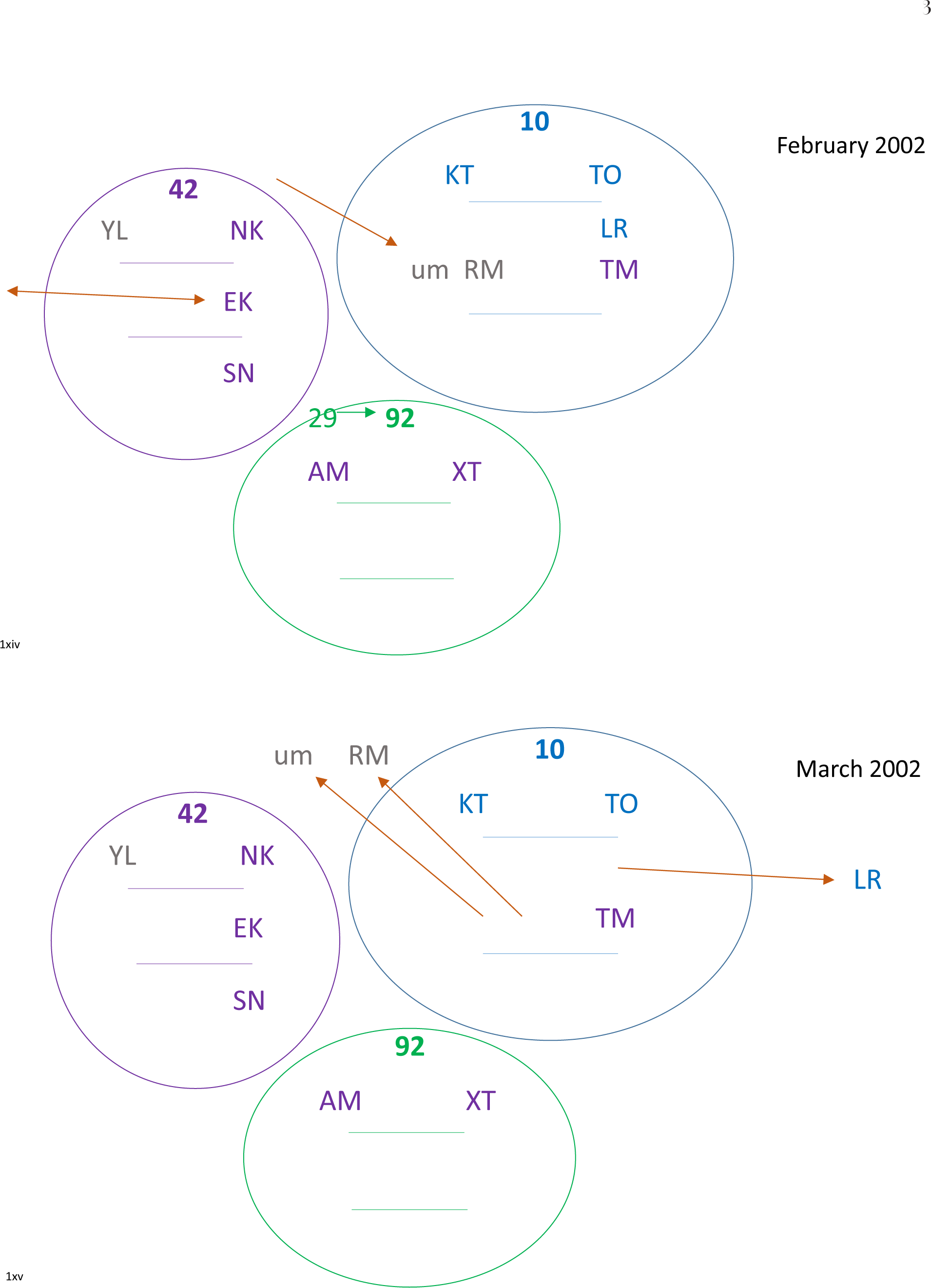
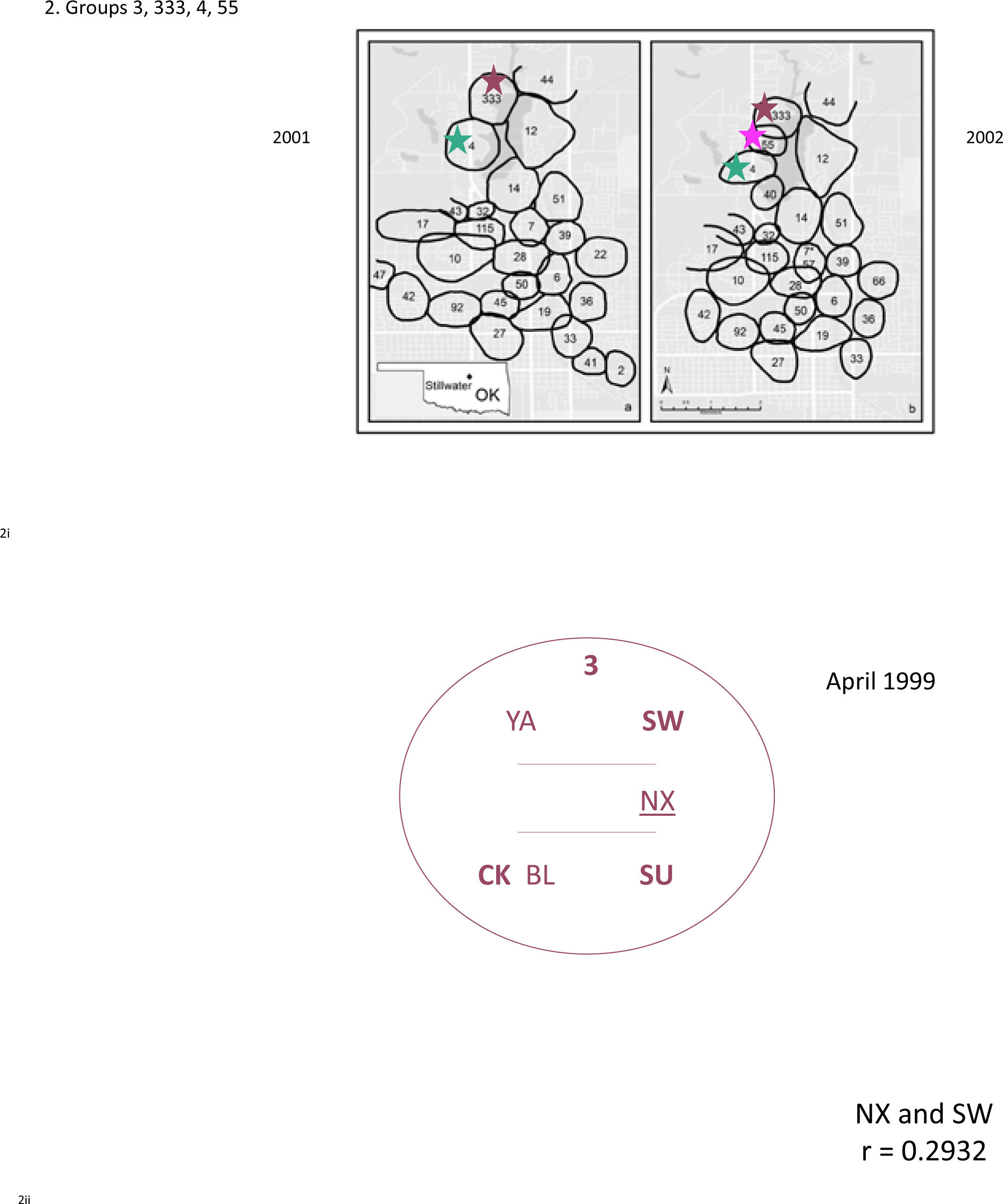
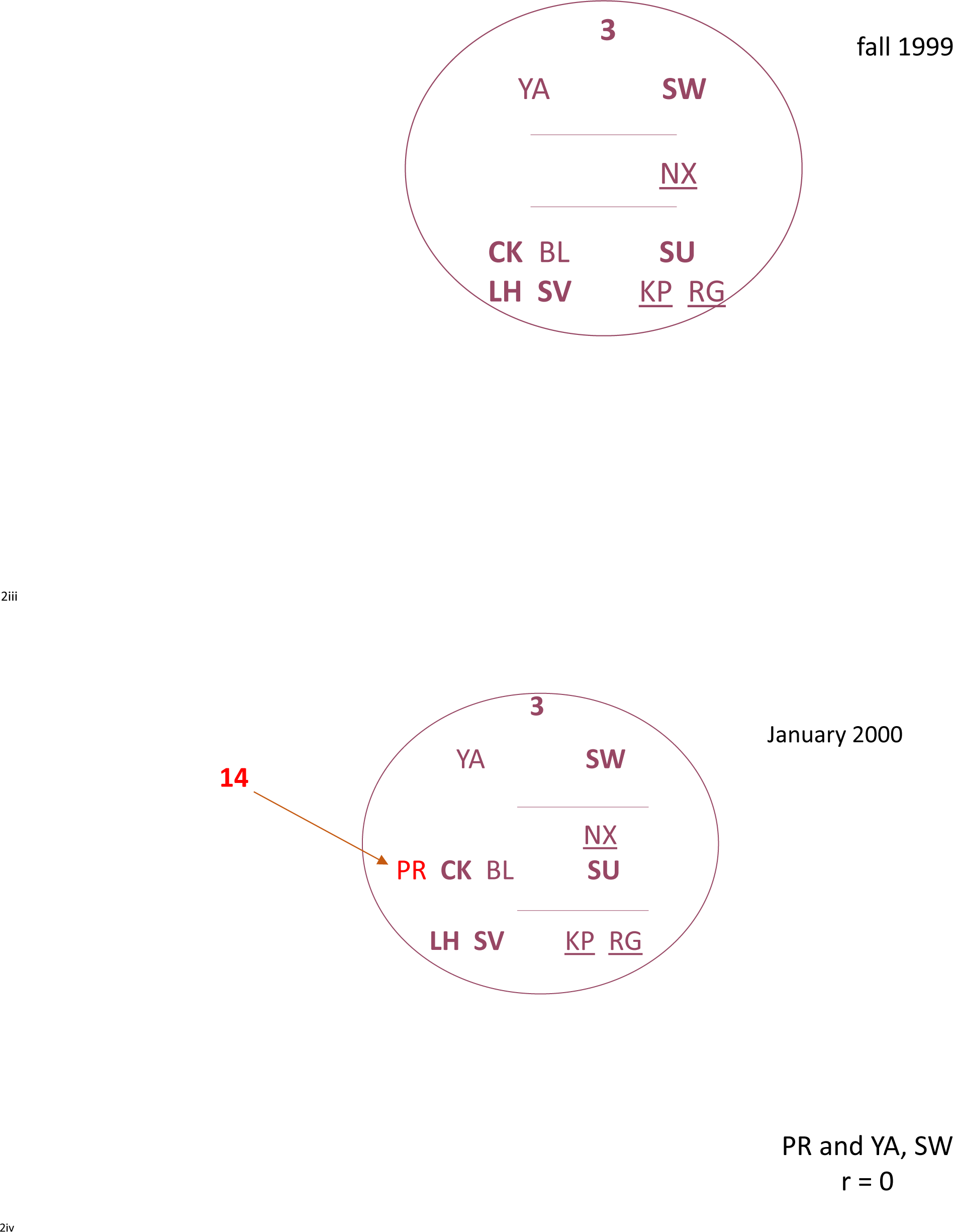
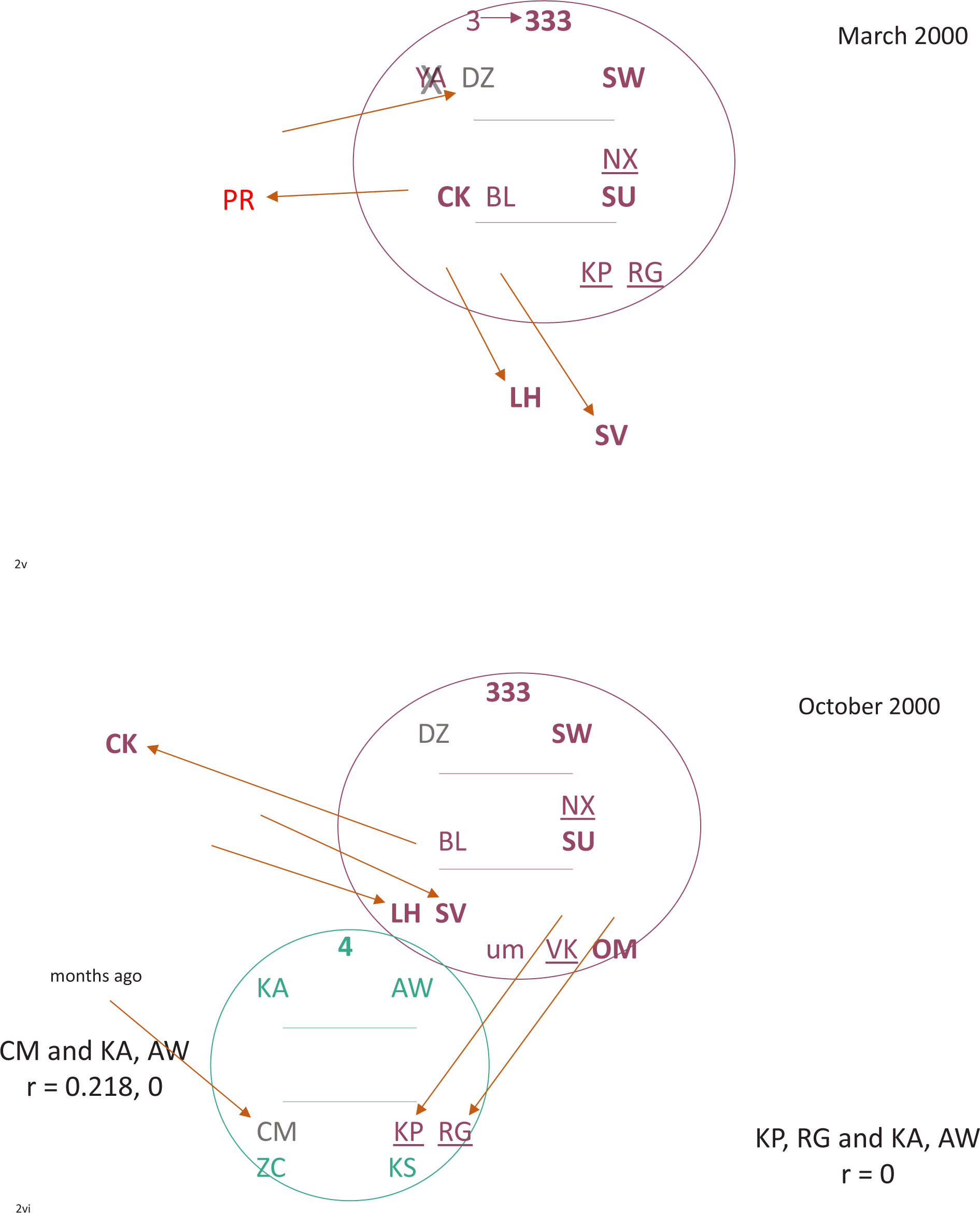
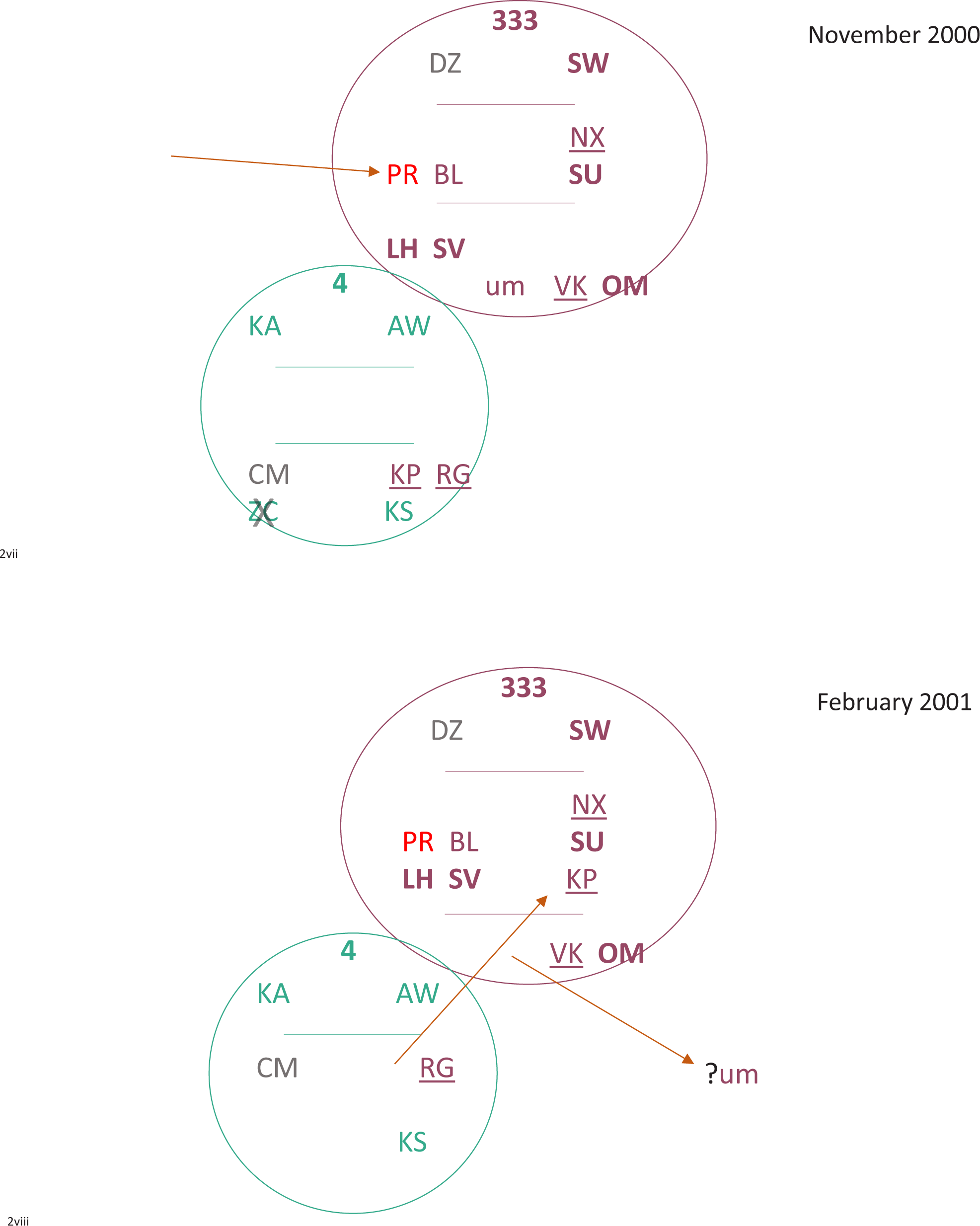
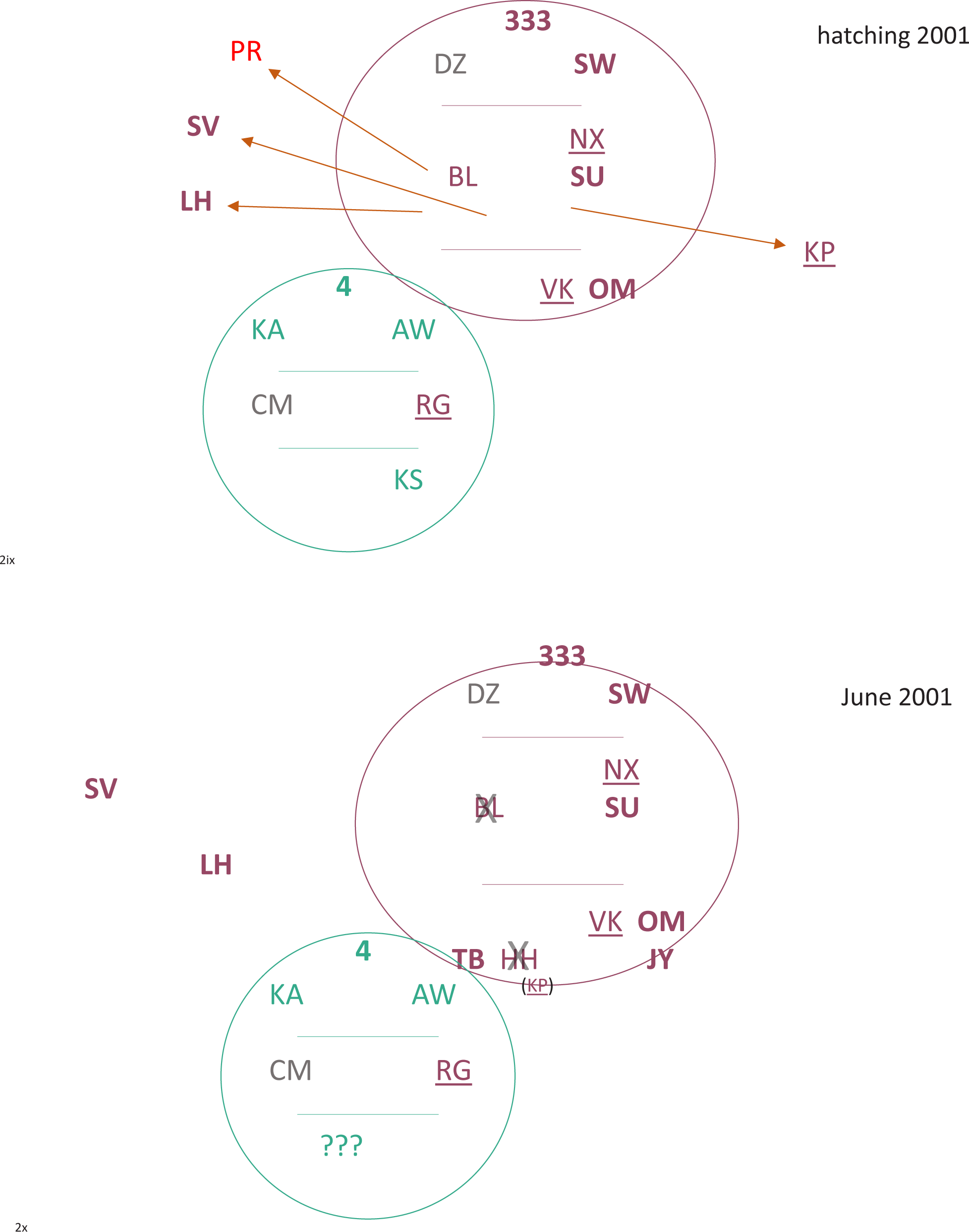
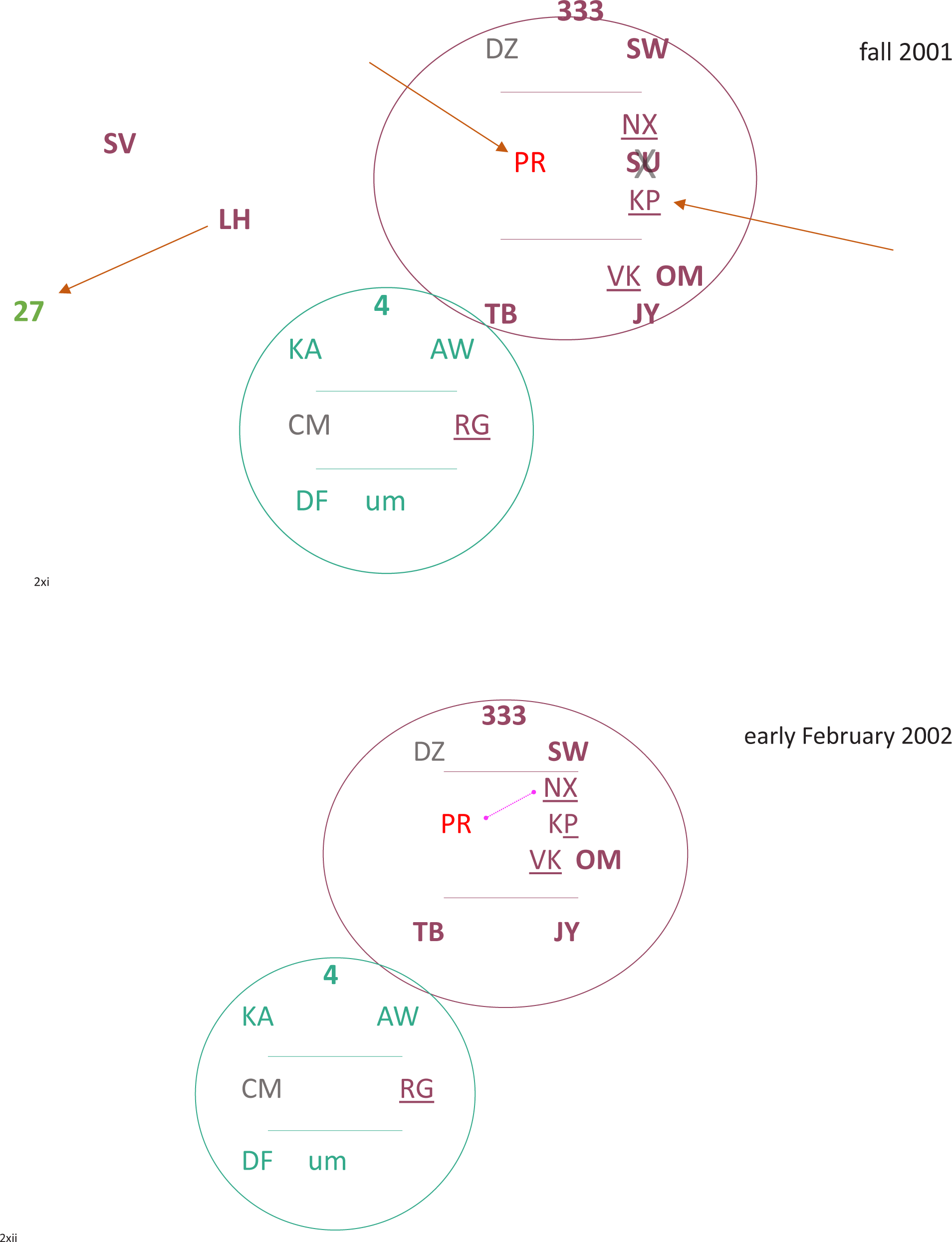
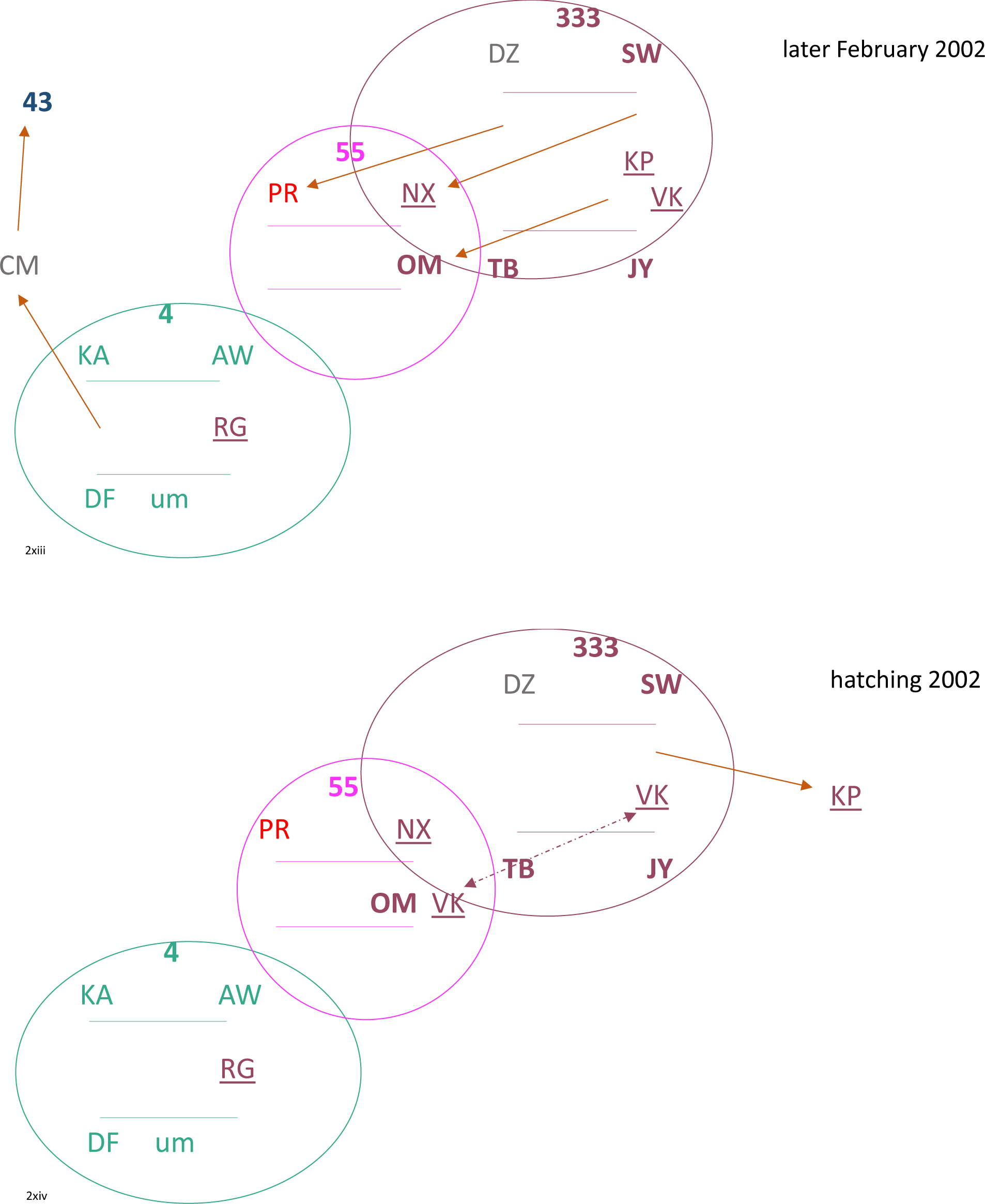
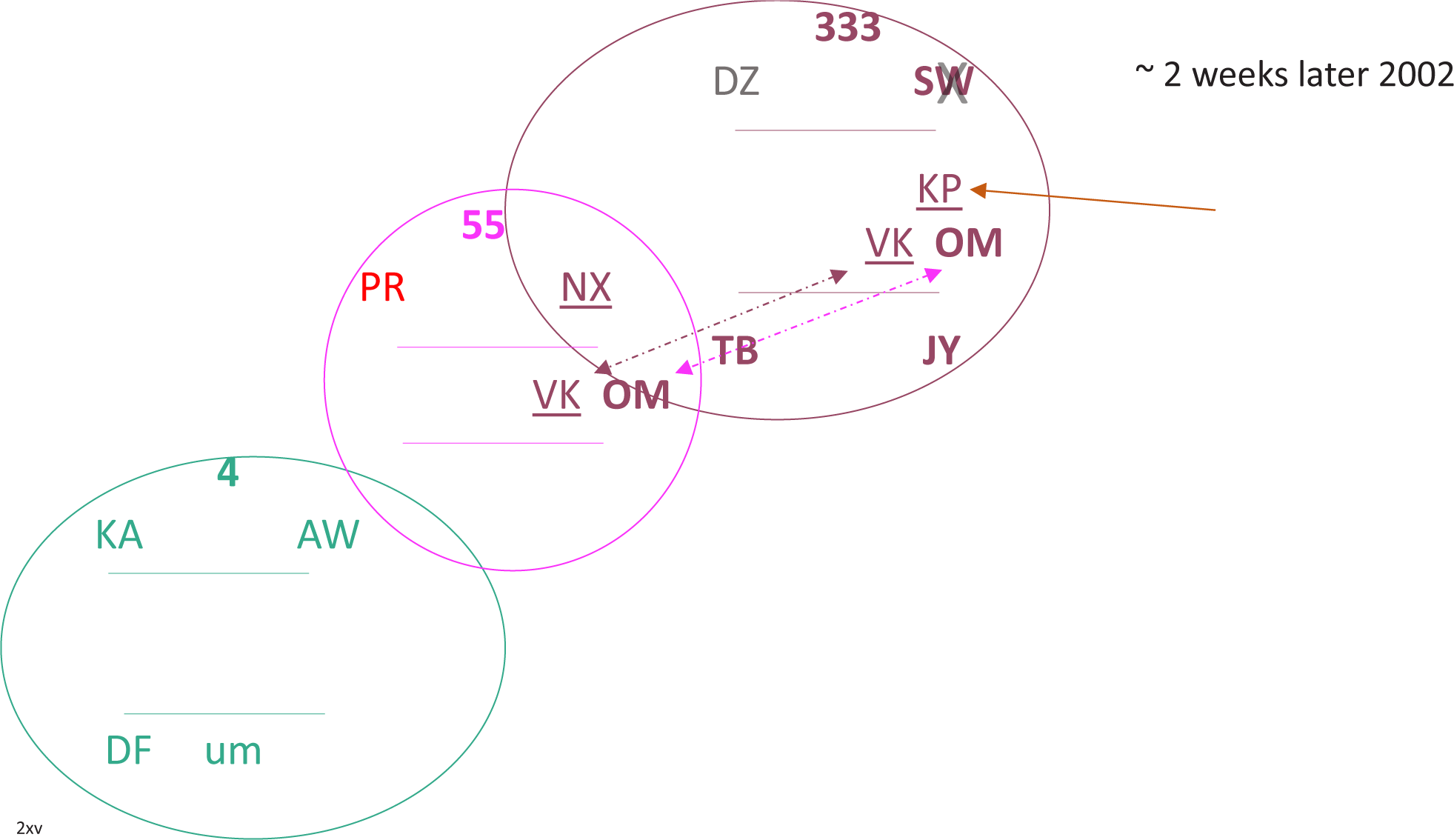

## APPENDIX D.

### Marked crows

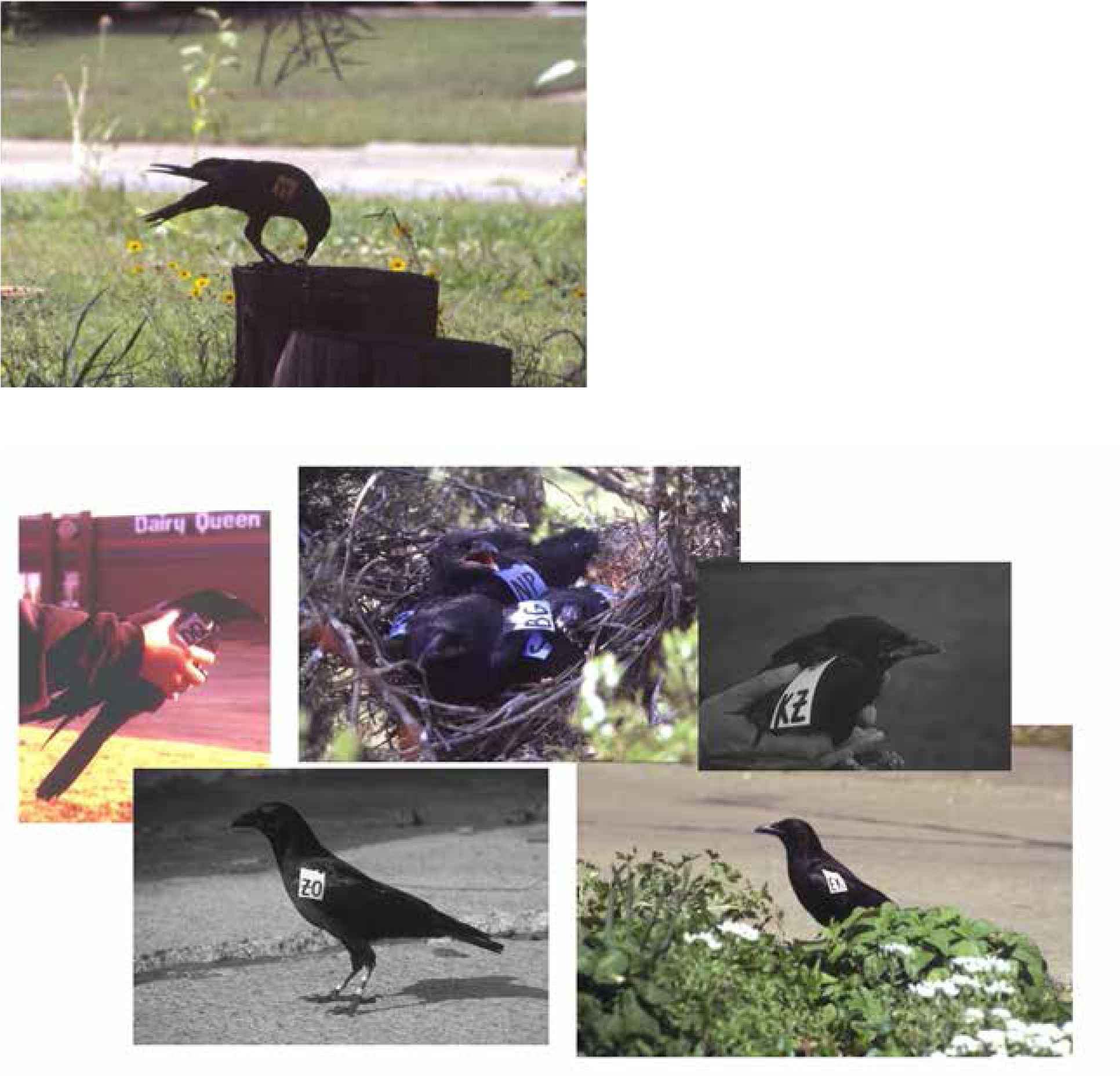

